# BioMapAI: Artificial Intelligence Multi-Omics Modeling of Myalgic Encephalomyelitis / Chronic Fatigue Syndrome

**DOI:** 10.1101/2024.06.24.600378

**Authors:** Ruoyun Xiong, Elizabeth Aiken, Ryan Caldwell, Suzanne D. Vernon, Lina Kozhaya, Courtney Gunter, Lucinda Bateman, Derya Unutmaz, Julia Oh

## Abstract

Myalgic Encephalomyelitis/Chronic Fatigue Syndrome (ME/CFS) is a chronic illness with a multifactorial etiology and heterogeneous symptomatology, posing major challenges for diagnosis and treatment. Here, we present BioMapAI, a supervised deep neural network trained on a four-year, longitudinal, multi-omics dataset from 249 participants, which integrates gut metagenomics, plasma metabolomics, immune cell profiling, blood laboratory data, and detailed clinical symptoms. By simultaneously modeling these diverse data types to predict clinical severity, BioMapAI identifies disease- and symptom-specific biomarkers and robustly classifies ME/CFS in both held-out and independent external cohorts. Using an explainable AI approach, we construct the first connectivity map spanning the microbiome, immune system, and plasma metabolome in health and ME/CFS, adjusted for age, gender, and additional clinical factors. This map uncovers disrupted associations between microbial metabolism (e.g., short-chain fatty acids, branched-chain amino acids, tryptophan, benzoate), plasma lipids and bile acids, and heightened inflammatory responses in mucosal and inflammatory T cell subsets (MAIT, γδT) secreting IFNγ and GzA. Overall, BioMapAI provides unprecedented systems-level insights into ME/CFS, refining existing hypotheses and hypothesizing new pathways associated to the disease’s heterogeneous symptoms.

## Introduction

Myalgic Encephalomyelitis/Chronic Fatigue Syndrome (ME/CFS) is a debilitating, multi-system illness that often persists for years or even decades and presents with substantial heterogeneity in clinical manifestations. Affecting an estimated 10 million individuals worldwide, ME/CFS is characterized by persistent fatigue, post-exertional malaise, multi-site pain, sleep disturbances, orthostatic intolerance, cognitive impairment, gastrointestinal symptoms, and other issues. This complexity not only hinders timely diagnosis but also poses significant challenges for effective treatment.^1,2,3^. The pathogenesis of ME/CFS is not well understood, with some triggers believed to include viral infections such as Epstein-Barr Virus (EBV)^4^, enteroviruses^5^ and SARS coronavirus^6^, in addition to bacterial infections and other causes^7^. As a chronic disease, ME/CFS can persist for years or even a lifetime, with each patient developing distinct illness patterns^1^. Hence, a single standardized approach to clinical care and symptom management is unlikely to suffice; instead, personalized, symptom-specific strategies may be necessary to effectively address the multifaceted nature of ME/CFS.

For ME/CFS and other chronic diseases such as cancer^8^, diabetes^9^, rheumatoid arthritis (RA)^10^, and long COVID^11,12^, this heterogeneity has been problematic to accommodate in research studies, leaving substantial knowledge and technical gaps^13^. The approach of most cohort studies is to focus on identifying one or two key disease indicators, such as HbA1C levels for diabetes^14,15^ or survival rates for cancer^16^, even with the advent of multi-‘omics. This approach has difficulty accommodating the highly multifactorial etiology and progression of most chronic diseases, with different patients exhibiting varying symptoms and disease markers^17^. To address this challenge, methods must link a more complex matrix of disease-associated outcomes with a range of ‘omics data types to enable precise targeting of biomarkers tailored to each patient’s specific symptoms.

In this study, we generated and assembled a longitudinal, multi-omics dataset from 153 ME/CFS patients and 96 age- and gender-matched healthy controls, encompassing gut metagenomics, plasma metabolomics, immune cell profiling (including activation and cytokine measures), blood labs, detailed clinical symptoms, and lifestyle surveys. To integrate these diverse data types with ME/CFS symptomatology, we developed BioMapAI, an explainable supervised deep neural network (DNN) that maps multi-omics profiles to a matrix of clinical symptoms. We aimed to: (1) identify novel disease biomarkers for ME/CFS, including those specifically tied to its heterogeneous symptomatology, and (2) map interactions among the microbiome, immune system, and metabolome rather than focusing on single or pairwise data types.

Using BioMapAI, we identified both disease- and symptom-specific biomarkers, reconstructed key clinical symptoms, and accurately classified ME/CFS in held-out and external cohorts. We then constructed a comprehensive multi-omics connectivity map that refines existing hypotheses and proposes new ones regarding microbial, metabolomic, and immune factors in ME/CFS. Critically, we accounted for confounders such as age and gender to contextualize the interplay among data types in health versus disease. For example, we observed that depletion of microbial short-chain fatty acids (e.g., butyrate) and branched-chain amino acids (BCAAs) in ME/CFS is linked to abnormal activation of mucosal and inflammatory immune cells (MAIT and γδT), which produce IFNγ and GzA—an altered dynamic correlated with worse perceived health and reduced social activity. Furthermore, microbial metabolites such as tryptophan and benzoate displayed fewer connections with plasma lipids in patients, an association that in turn tracked with fatigue, emotional dysregulation, and sleep disturbances.

To our knowledge, this dataset is among the most comprehensive multi-omics resources assembled for ME/CFS (including other complex chronic diseases). We further introduce an innovative AI approach that begins to address the multifaceted nature of this chronic disease, generating new hypotheses for host–microbiome interactions in both health and ME/CFS. Given the recognized parallels in both etiology and clinical presentation between ME/CFS and long COVID^11,12^, studying ME/CFS can offer broader insights into the pathophysiology of post-viral syndromes. More generally, our AI-driven framework may prove valuable for other complex conditions where symptom variability cannot be fully captured by a single data type.

## Results

### Cohort Overview

We tracked 249 participants over 3-4 years, including 153 ME/CFS patients (75 ‘short-term’ with disease symptoms < 4 years and 78 ‘long-term’ with disease symptoms > 10 years) and 96 healthy controls (Fig 1A; Supplemental Table 1). The cohort is 68% female and 32% male, aligning with the epidemiological data showing that women are 3-4 times more likely to develop ME/CFS^18,19^. Participants ranged in age from 19 to 68 years with body mass indexes (BMI) from 16 to 43 kg/m². Throughout the study, we collected detailed clinical metadata, blood samples, and fecal samples. In total, 1471 biological samples were collected across all participants at 515 timepoints (Methods, Supplemental Figure 1A, Supplemental Table 1).

**Figure 1:**
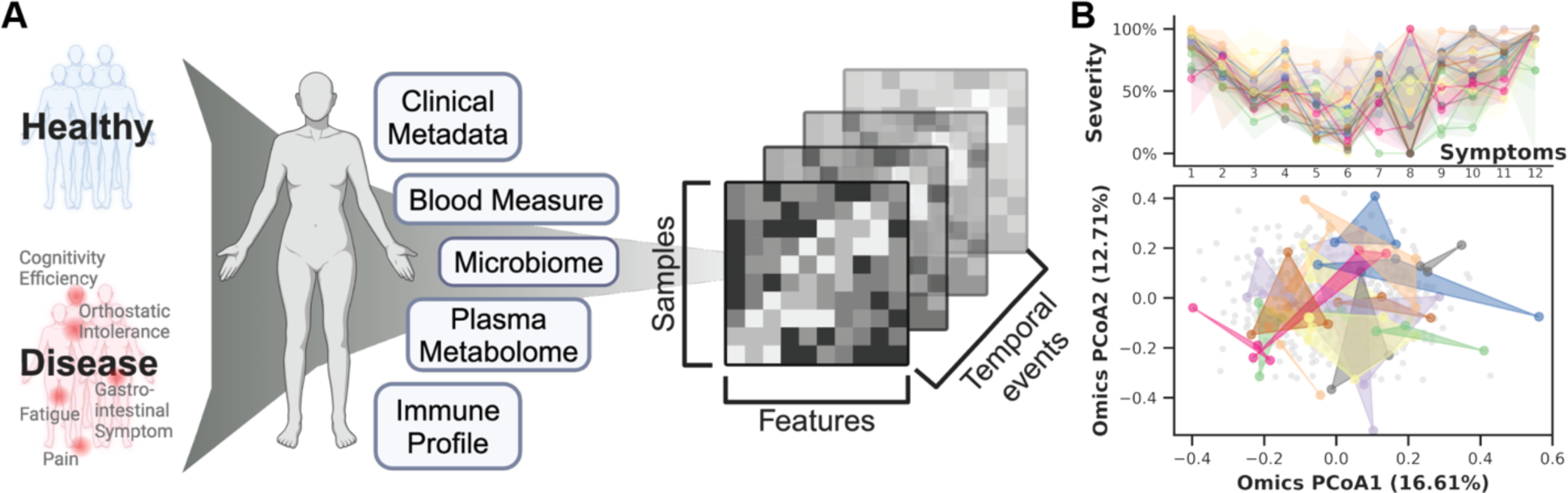
Cohort Summary and Heterogeneity of ME/CFS. A) Cohort Design and ‘Omics Profiling. 96 healthy donors and 153 ME/CFS patients were followed over 3-4 years with yearly sampling. Clinical metadata including lifestyle and dietary surveys, blood clinical laboratory measures (N=503), gut microbiome (N=479), plasma metabolome (N=414), and immune profiles (N=489) were collected (Supplemental Table 1 and Supplemental Figure 1A). **B) Heterogeneity and Non-Linear Progression of ME/CFS in Symptom Severity and ‘Omics Profiles.** This section highlights variability in symptom severity (top) and ‘omics profiles (bottom) for 20 representative ME/CFS patients over 3–4 time points. **Top,** Symptom severity is shown for 12 major clinical symptoms (x-axis, with each column representing one symptom) against severity scores (scaled from 0% (no symptom) to 100% (most severe), y-axis) for each patient (each represented by a distinct color). Lines indicate average severity, and shaded areas represent the severity range across time points (controls shown in Supplemental Figure 1B). Here, we observed a lack of consistent temporal patterns for ME/CFS symptomatology, indicated by the widespread shaded areas, and significant heterogeneity over time (Supplemental Figure 1F–G). Notably, among the 12 symptoms, trends differed: fatigue (Symptom 1) remains consistently severe over years, whereas emotional dysregulation (Symptom 8) exhibit notable variability and instability over time. **Bottom**, PCoA of integrated ‘omics data with color dots matching patient timepoints in the symptom plot and grey dots representing the entire cohort. Again, the spread and overlap of the colored space reflect the diversity in ‘omics signatures vs. the more consistent pattern typical of controls (Supplemental Figure 1C). **Abbreviations:** ME/CFS, Myalgic Encephalomyelitis/Chronic Fatigue Syndrome; PCoA, Principal Coordinates Analysis. **Supporting Materials:** Supplemental Table 1, Supplemental Figure 1.

Blood samples were 1) sent for clinical testing at Quest Laboratory (48 features measured, N=503 samples), 2) fractionated into peripheral blood mononuclear cells (PBMCs), which were examined via flow cytometry, yielding data on 443 immune cells and cytokines (N=489), 3) plasma and serum, for untargeted liquid chromatography with tandem mass spectrometry (LC-MS/MS), identifying 958 metabolites (N=414). Detailed demographic documentation and questionnaires covering medication use, medical history, and key ME/CFS symptoms were collected (Methods). Finally, whole genome shotgun metagenomic sequencing of stool samples (N=479) produced an average of 12,302,079 high-quality, classifiable reads per sample, detailing gut microbiome composition (1293 species detected) and KEGG gene function (9993 genes reconstructed).

### Heterogeneity and Non-linear Progression of ME/CFS

First, we demonstrated the phenotypic complexity and heterogeneity of ME/CFS. Collaborating with clinical experts, we consolidated detailed questionnaires and clinical metadata foundational to diagnosing ME/CFS, into twelve essential clinical scores (Methods). These scores covered core symptoms including physical and mental health, fatigue, pain levels, cognitive efficiency, sleep disturbances, orthostatic intolerance, and gastrointestinal issues (Supplemental Table 1).

While healthy individuals consistently presented low symptom scores (Supplemental Figure 1D, 1F), ME/CFS patients exhibited significant variability in symptom severity, with each individual showing different predominant symptoms (Figure 1B, Supplemental Figure 1C, 1G). Principal coordinates analysis (PCoA) of the ‘omics matrices highlighted the difficulty in distinguishing patients from controls, emphasizing the complex symptomatology of ME/CFS and the challenges in developing predictive models (Supplemental Figure 1E). Additionally, over time, in contrast to the stable patterns typical of healthy individuals (Supplemental Figure 1B), ME/CFS patients demonstrated distinctly varied patterns each year, as evidenced by the diversity in symptom severity and noticeable separation on the ‘omics PCoA (Figure 1B, Supplemental Figure 1C). Despite employing multiple longitudinal models (Methods), we found no consistent temporal signals, confirming the non-linear progression of ME/CFS.

This individualized, multifaceted, and dynamic nature of ME/CFS that intensifies with disease progression necessitates new approaches that extend beyond simple disease versus control comparisons. Here, we created and implemented an AI-driven model that integrates the multi-’omics profiles to learn host phenotypes. This allowed us not only to develop an accurate disease classifier, but more importantly, to identify specific biomarker sets for each clinical symptom as well as unique interaction networks that differed between patients and controls.

### BioMapAI, an Explainable Neural Network Connecting ‘Omics to Multi-Type Outcomes

ME/CFS research is hindered by the complexity of its clinical phenotypes and biological measurements, which are highly individualized. To associate multi-’omics data with clinical symptoms, a model must accommodate the learning of multiple different outcomes within a single framework. However, traditional machine learning models are generally designed to predict a single categorical outcome or continuous variable^20,21,22^. This simplified disease classification and conventional biomarker identification typically fails to encapsulate the heterogeneity of complex diseases^23,24^. Our goal was to integrate multi-‘omics data with clinical symptoms into a single model, which would enable a direct comparison of the predictive value of different ‘omics datasets and the identification of symptom-specific biomarkers within a unified framework.

We developed an AI-powered multi-’omics framework, BioMapAI, a fully connected deep neural network that inputs ‘omics matrices (*X*), and outputs a mixed-type outcome matrix (*Y*), thereby mapping multiple ‘omics features to multiple clinical indicators (Figure 2A). By assigning tailored loss functions for each output to each output based on its data type (See Methods), BioMapAI aims to comprehensively learn every *y* (i.e., each of the 12 continuous or categorical clinical scores in this study), using the ‘omics data inputs. Between the input layer *X* and the output layer *Y* = [*y*_1_, *y*_2_, …, *y*_*n*_], the model consists of two shared hidden layers (*Z*^1^ with 64 nodes, and *Z*^2^ with 32 nodes) for general pattern learning, followed by a parallel hidden layer (*Z*^3^ = [*z*_1_^3^, *z*_2_^3^, …, *z*_n_^3^]), with sub-layers (*z*_n_^3^, each with 8 nodes) tailored for each outcome (*y*_n_), to capture outcome-specific patterns (Figure 2A). This unique architecture – two shared and one specific hidden layer – allows the model to capture both general and output-specific patterns. This model is made 1) explainable by incorporating a SHAP (SHapley Additive exPlanations) explainer, which quantifies the feature importance of each predictions, providing both local (symptom-level) and global (disease-level) interpretability, and 2) flexible by automatically finding appropriate learning goals and loss functions for each type of outcomes (without need of format refinement), facilitating BioMapAI’s potential adaptability to broader research applications.

**Figure 2:**
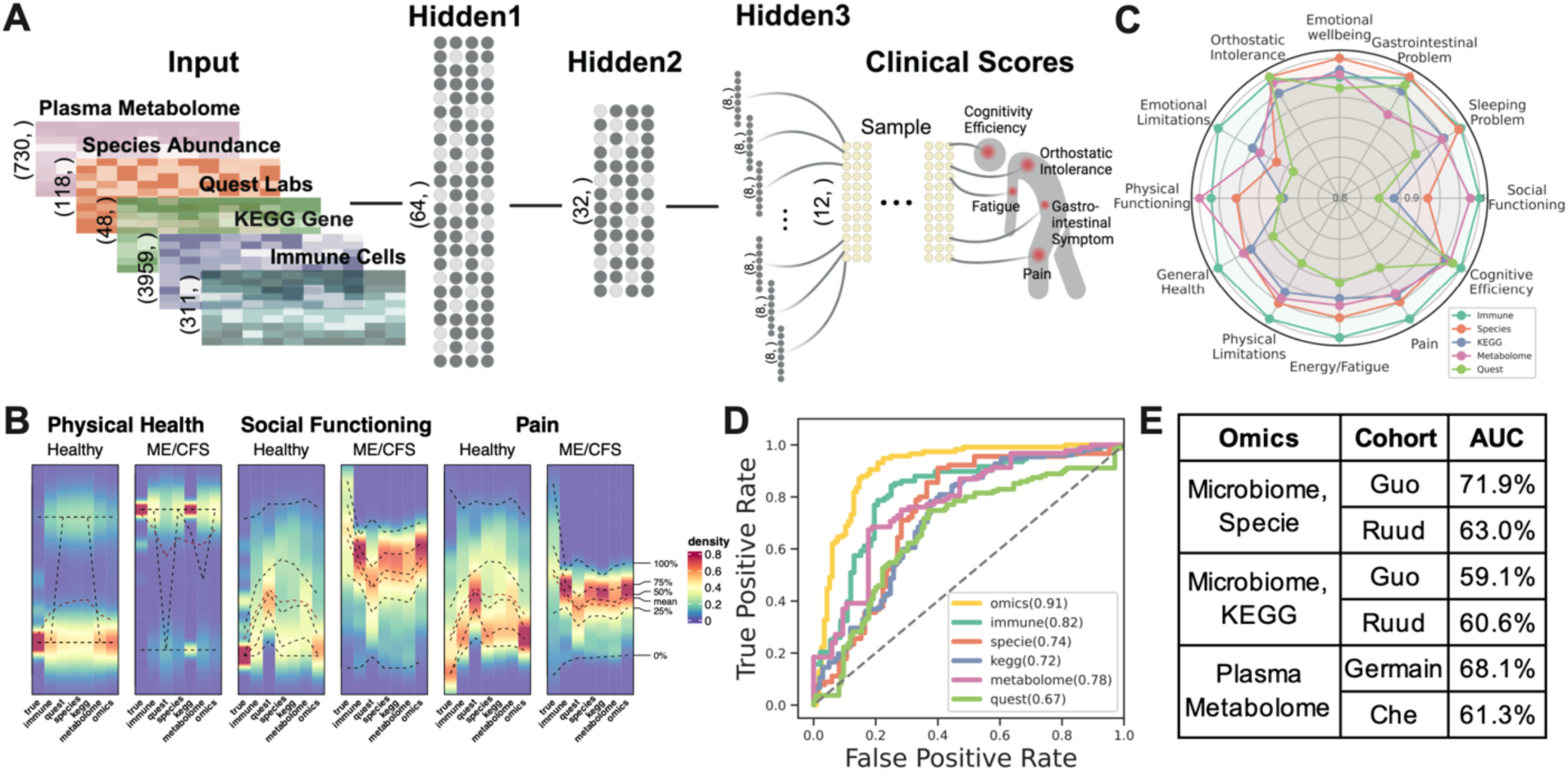
BioMapAI’s Model Structure and Performance. A) Structure of BioMapAI. BioMapAI is a fully connected deep neural network comprised of an input layer (*X*), a normalization layer (not shown), three sequential hidden layers (*Z*^1^, *Z*^2^, *Z*^3^), and one output layer (*Y*). Hidden layer 1 (*Z*^1^, 64 nodes) and hidden layer 2 (*Z*^2^, 32 nodes), both feature a dropout ratio of 50% to prevent overfitting (visually represented by dark and light gray nodes). Hidden layer 3 has 12 parallel sub-layers each with 8 nodes (*Z*^3^ = [*z*_1_^3^, *z*_2_^3^, …, *z*_12_^3^]) to learn 12 objects in the output layer (*Y* = [*y*_1_, *y*_2_, …, *y*_12_]) representing key clinical symptoms of ME/CFS. **B) True vs. Predicted Clinical Scores highlight BioMapAI’s accuracy.** Three example density maps (full set, Supplemental Figure 2A) compare the true score, *y* (Column 1) against BioMapAI’s predictions generated from different ‘omics profiles - ŷ_*immune*_, ŷ_*speices*_,ŷ_*KEGG*_, ŷ_*metabolome*_, ŷ_*quest*_, ŷ_*omics*_ (Columns 2-7). Y-axis represents the diversity calculated by kernel density estimation (KDE), which is a smoothed estimate of the distribution of the symptom severity along the x-axis for each omics. Color gradient from blue (lower density) to red (higher density) illustrates the occurrence frequency (e.g., true scores for ∼100% of healthy controls’ physical health ∼ 0 = red), with dashed lines indicating key statistical percentiles (100%, 75%, 50%, 25%, and 0%). Note that model’s predicted scores a preserve differences between healthy controls and patients for these three examples, irrespective of ‘omics type. **C) ‘Omics’ Strengths in Symptom Prediction.** Radar plot shows BioMapAI’s performance in predicting the 12 clinical outcomes for each ‘omics datatype. Each of the 12 axes represents a clinical score output (*Y* = [*y*_1_, *y*_2_, …, *y*_12_]), with five colors denoting the ‘omics datasets used for model training. The spread of each color along an axis reflects the 1 - normalized mean square error (MSE) (Supplemental Table 2) between the actual, *y*, and the predicted, *y*2, outputs, illustrating the predictive strength or weakness of each ‘omics for specific clinical scores. The radial scale ranges from 0.8 (center) to 1.0 (outer circle), where values closer to the outer edge correspond to lower MSE and better predictions. For instance, species abundance predicted gastrointestinal, emotional, and sleep issues effectively, while the immune profile was broadly accurate across most scores. **D) BioMapAI’s Performance in Healthy vs. Disease Classification (10-Fold Cross Validation).** ROC curves show BioMapAI’s performance in disease classification using each ‘omics dataset separately or combined (‘Omics’), with the AUC in parentheses showing prediction accuracy (full report in Supplemental Table 3, held out data ROC in Supplemental Figure 2E). **E) Validation of BioMapAI with External Cohorts.** External cohorts with microbiome data (Guo et al.^25^, Ruud et al.^26^) and metabolome data (Germain et al.^27^, Che et al.^32^) were used to test BioMapAI’s model, underscoring its generalizability (detailed classification matrix, Supplemental Table 4). **Abbreviations:** KEGG, Kyoto Encyclopedia of Genes and Genomes; ‘Omics’ refers to the combined multi-‘omics matrix; MSE, Mean Square Error; ROC curve, Receiver Operating Characteristic curve; AUC, Area Under the Curve; *y*, True Score; *y*2, Predicted Score. **Supporting Materials:** Supplemental Tables 2-4, Supplemental Figures 1-2.

### BioMapAI Reconstructed Clinical Symptoms and Demonstrated Robust Capability to Classify ME/CFS from Healthy Controls

BioMapAI is a supervised deep learning AI framework that connects a biological ‘omics matrix to multiple phenotypic outputs. Here, we trained and validated it on our ME/CFS dataset, employing a ten-fold cross-validation. Additionally, 10% of the data was held out as an independent validation set, separate from the cross-validation process, to assess the model’s generalizability (Methods). This trained model, nicknamed DeepMECFS for the ME/CFS community, was able to represent the structure of diverse clinical symptom score types and discriminated between healthy individuals and patients (Figure 2, Supplemental Figure 2, Supplemental Table 2-3). For example, it effectively differentiated the physical health scores, where patients exhibited more severe conditions compared to healthy controls (category datatype 4 vs. 0, respectively, Figure 2B, Supplemental Table 2) and pain scores (continuous datatype ranging from 1(highest)-0(lowest), mean 0.52±0.24 vs. 0.11±0.12 for patients vs. controls). Though compressing some inherent variance, BioMapAI accurately reconstructed key statistical measures such as the mean and interquartile range (25%-75%), and highlighted the distinctions between healthy and disease. (Figure 2B, Supplemental Figure 2A-B, Supplemental Table 2).

To determine the accuracy of BioMapAI’s reconstructed clinical scores, we compared their ability to discriminate ME/CFS patients from controls with the original clinical scores. We used one additional fully connected layer to regress the 12 predicted clinical scores Ŷ(12,) into a binary outcome of patient vs. control ŷ_(1,)_. Because the diagnosis of ME/CFS relies on clinical interpretation of key symptoms (i.e., the original clinical scores), the original clinical scores have near-perfect accuracy in classification, as expected (AUC, Area Under the Curve >99%, Supplemental Figure 2C). BioMapAI’s predicted scores achieved a 91% AUC in distinguishing disease from healthy controls as evaluated through 10-fold cross-validation (Figure 2D, Supplemental Figure 2D). To benchmark its performance, we compared it with four machine learning models - generalized linear model with elastic net regularization (Glmnet), Glmnet with interaction terms, support vector machine (SVM), and gradient boosting (GDBT) – and a deep learning model (DNN) with two fully connected layers but without the third “spread-out” hidden layer (Supplemental Table 3). In terms of the 10-fold cross-validation for disease classification, BioMapAI, DNN, and Glmnet performed comparably well overall. BioMapAI showed slightly better performance with the full ‘omics dataset (AUC = 91.5%) and immune data (81.8%), while Glmnet outperformed in metabolome (79.0%) and questionnaire data (72.5%).

BioMapAI demonstrated robust performance with unseen data, as validated on held-out cohort data (Supplemental Figure 2E, Table 3) and independent, previously published ME/CFS cohorts (Figure 2E, Supplemental Table 4). In the held-out validation, it outperformed in most ‘omics datasets, including ‘omics altogether (AUC=82.3%), immune (78.5%), KEGG (69.1%), species (71.5%), and metabolome (76.4%), while Glmnet excelled in Quest data (74.8%). Public datasets included two microbiome cohorts, Guo, Cheng et al., 2023 (US)^25^ and Raijmakers, Ruud et al., 2020 (Netherlands)^26^ and two metabolome cohorts, Germain, Arnaud et al., 2022 (US)^27^ and Che, Xiaoyu et al., 2022 (US)^28^. Despite the challenges of validating traditional microbiome and metabolite ML models using external cohorts – often having technical (e.g., metabolomic features only overlapped by 79% and 19% for the two studies, respectively) and clinical differences^29,30,31^, BioMapAI demonstrated good performance and outperformed other models (Figure 2E, Supplemental Table 4). While BioMapAI’s accuracy using these external datasets was lower, its improved performance highlights the value of incorporating clinical symptoms into a predictive model, demonstrating that connecting ‘omics features to clinical symptoms improves disease classification.

### ‘Omics’ Strengths Varied in Symptom Prediction; Immune is the Most Predictive

One innovation of BioMapAI is its ability to leverage different ‘omics data to predict individual clinical scores in addition to disease vs. healthy classification. We evaluated the predictive accuracy by calculating the mean squared error between actual (*y*) and predicted (*y*2) scores and observed that the different ‘omics showed varying strengths in predicting clinical scores (Figure 2C), likely due in part to the wide differences in dimensionality specific to each datatype. Immune profiling consistently had the highest ability to forecast a wide range of symptoms, including pain, fatigue, orthostatic intolerance, and general health perception, underscoring the immune system’s crucial role in health regulation. In contrast, blood measurements demonstrated limited predictive ability, except for cognitive efficiency, likely owing to their limited focus on 48 specific blood bioactives. Plasma metabolomics, which encompasses nearly a thousand measurements, performed significantly better with notable correlations with facets of physical health and social activity. These findings corroborate published metabolites and mortality^32,33^, longevity^34,35^, cognitive function^36^, and social interactions^37,38,39^. Microbiome profiles surpassed other ‘omics in predicting gastrointestinal abnormalities (as anticipated^40,41^), emotional well-being, and sleep disturbances, supporting recently established links in gut-brain health^42,43,44^.

### BioMapAI is Explainable, Identifying Disease- and Symptom-Specific Biomarkers

Deep learning (DL) models are often referred to as ‘black box’, with limited ability to identify and evaluate specific features that influence the model’s predictions. BioMapAI is made explainable by incorporating SHAP values, which quantify how each feature influenced the model’s predictions. BioMapAI’s architecture – two shared layers (*Z*^1^ and *Z*^2^) for general disease pattern learning and one parallel layer for each clinical score (*Z*^3^ = [*z*_1_^3^, *z*_2_^3^, …, *z*_12_^3^]) – allowed us to identify both disease-specific biomarkers, which are shared across symptoms and models (Supplemental Figure 3, Supplemental Table 5), and symptom-specific biomarkers, which are tailored to each clinical symptom (Figure 3, Supplemental Figure 4-5, Supplemental Table 6).

**Figure 3:**
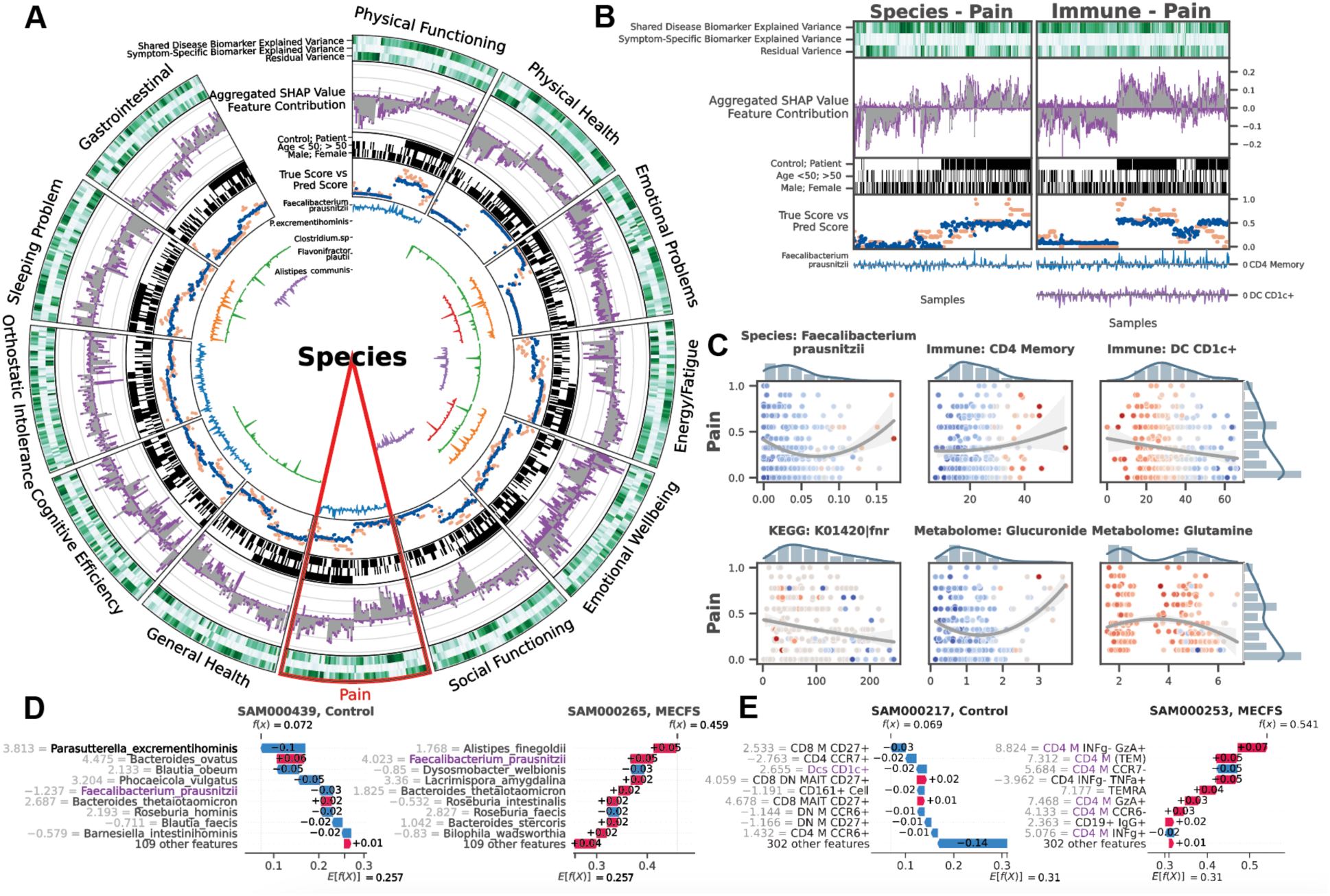
BioMapAI Identifies both Disease- and Symptom-Specific Biomarkers. For Symptom-Specific Biomarkers, A) Circularized Diagram of Species Model with B) Zoomed Segment for Pain. Each circular panel illustrates how the model predicts each of the 12 symptom-specific biomarkers derived from one type of ‘omics data (all datatypes shown in Supplemental Figure 4). The x-axis for each panel represents an individual’s values for each of the following contributors to the model’s performance (from top to bottom): *1. Variance Explained by Biomarker Categories:* Gradients of dark green (100%) to white (0%) show variance explained by the model. For many biomarkers, disease-specific biomarkers account for the greatest proportion of variance, and symptom-specific biomarkers provide additional tailored explanations, with residual accounting for the remaining variance; *2. Aggregated SHAP Values* quantify the contribution of each feature to the model’s predictions, with disease-specific biomarkers in grey and symptom-specific in purple. *3. Demography and Cohort Classification:* cohort (controls, white vs. patients, black); age <50 (white) vs. >50 years old (black); sex (male, white vs. female, black); *4. True vs. Predicted Scores* show BioMapAI’s predictive performance at the individual sample level, with true in blue and model-predicted scores in orange; *5.Examples of Symptom-Specific Biomarkers:* Line graphs show the contribution of select symptom-specific biomarkers to the model across individuals, e.g., 5 gut species in A). In B), the three features most specific to the pain model include gut microbe *F. prausnitzii,* CD4 memory T, and DC CD1c+ cells. Peaks above 0 (middle line) indicate a positive contribution and below 0 for a negative contribution. For example, the mixed positive and negative contribution peaks of *F. prausnitzii* indicated a biphasic contribution to pain intensity. Disease-Specific Biomarkers are shown in Supplemental Figure 3. **C) Different Correlation Patterns of Biomarkers to Symptoms:** For pain (other symptoms in Supplemental Figure 5), correlation analysis of raw abundance (x-axis) of each biomarker with pain score (y-axis) show monotonic (e.g., CD4 memory and DC CD1c+ markers), biphasic (microbial and metabolomic markers), or sparse (KEGG genes) contribution patterns for those features. Dots represent an individual color-coded to SHAP value, where the color spectrum indicates negative (blue) to neutral (grey) to positive (red) contributions to pain prediction. Superimposed trend lines with shaded error bands represents the predicted correlation trends between biomarkers and pain intensity. Adjacent bar plots represent the data distribution. **D-E) Examples of Pain-Specific Biomarkers’ Contributions.** SHAP waterfall plots (colors corresponding to gradient in C) illustrate the contribution of individual features to a model’s predictive output. The top 10 features for two pairs of controls and patients are shown here, illustrating the species and the immune model (additional examples in Supplemental Figure 4A). The contribution of each feature is shown as a step (SHAP values provided adjacent), and the cumulative effect of all the steps provides the final prediction value, *E*[*f*(*X*)]. Our example of *F. prausnitzii* exhibits a protective role (negative SHAP) in controls but exacerbates pain (positive SHAP) in patients – consistent with the biphasic relationship observed in C). As a second example, all CD4 memory cells in this model have positive SHAP values, reinforcing the positive monotonic relationship with pain severity observed in C). Conversely, DC CD1c+ cells contribute negatively and thus may have a protective role. *Note, the reported biomarkers were calculated using the entire dataset and were not validated on held-out data. **Abbreviation:** SHAP, SHapley Additive exPlanations; DNN, Deep Neuron Network; GBDT, Gradient Boosting Decision Tree; KEGG, Kyoto Encyclopedia of Genes and Genomes. **Supporting Materials:** Supplemental Table 5-6, Supplemental Figure 3-5.

Disease-specific biomarkers are important features across symptoms and models (Methods, Supplemental Figure 3). Increased B cells (CD19+CD3-), CCR6+ CD8 memory T cells (mCD8+CCR6+CXCR3-), and CD4 naïve T cells (nCD4+FOXP3+) in patients were associated with most symptoms, suggesting a potentially broad dysregulation of the adaptive immune response. The species model highlighted the importance of *Dysosmobacteria welbionis*, a gut microbe previously reported in obesity and diabetes, with a role in bile acid and butyrate metabolism^45,46^. The metabolome model categorized increased levels of glycodeoxycholate 3-sulfate, a bile acid, and decreased vanillylmandelate (VMA), a catecholamine breakdown product^47^. These features shared for all symptoms were consistently validated across ML and DL models, demonstrating the efficacy of BioMapAI (Supplemental Table 5).

More uniquely, BioMapAI linked ‘omics profiles to clinical symptoms and thus enabled the identification of symptom-specific biomarkers (Figure 3A). Certain ‘omics data, like species-gastrointestinal and immune-pain associations, were especially effective in predicting specific clinical phenotypes (Figure 2C). Utilizing SHAP, BioMapAI identified distinct sets of biomarkers for each symptom (Supplemental Table 6, Supplemental Figure 5). We found that while disease-specific biomarkers accounted for a substantial portion of the variance, symptom-specific biomarkers crucially refined the predictions, aligned predicted scores – consistently across age and gender – more closely with actual values (Figure 3A-B, Supplemental Figure 4B-D). For example, in the case of pain, CD4 memory and CD1c+ dendritic cells (DC) were particularly important features, and *Faecalibacterium prausnitzii* was also uniquely associated, with varying impact across individual (Figure 3B). Similar to pain, each clinical score in ME/CFS was characterized by its unique ‘omics features, distinct from those common across other symptoms (Supplemental Table 6).

In addition, we observed a spectrum of interaction types (linear, biphasic, and dispersed) extending beyond conventional linear interactions, underscoring the heterogeneity inherent in ME/CFS (Figure 3C). High-abundance species and immune cells often had a biphasic relationship with symptoms, showing dual effects, while low-abundance species and metabolites displayed a linear relationship with positive or negative associations with clinical scores (Supplemental Figure 5).

An example of a relatively straightforward monotonic (linear) relationship was observed between CD4 memory (CD4 M) cells, CD1c+ DCs and pain, with positive associations of CD4 M cells to pain intensity severity. Conversely, CD1c+ DCs had negative associations to pain severity in both patients and control (Figure 3C, E). These variations suggest alterations in inflammatory responses and specific pathogenic processes in ME/CFS, which may be virally triggered and is marked by prolonged infection symptoms. Many microbial biomarkers demonstrated linear contributions to symptoms, evidenced by numerous negative peaks indicating a positive association in symptom severity (Figure 3A). For example, *Dysosmobacteria welbionis*, a disease-specific biomarker, was associated with more severe sleeping and gastrointestinal issues (Supplemental Figure 3), whereas *Clostridium sp.* and *Alistipes communis* were associated with less severe scores (Figure 3A, Supplemental Figure 5B).

A more complex, biphasic relationship was observed in the association of *Faecalibacterium prausnitzii* with pain, whose saddle curve (Figure 3C) had a mixture of positive and negative contribution peaks (Figure 3B), which means that either abnormally low and high relative abundances could be associated with pain severity. In disease, *F. prausnitzii* was associated with higher pain scores, while in healthy individuals, it was associated with lower pain scores (Figure 3D). Notably, *F. prausnitzii* was identified as a biomarker in several ME/CFS cohorts^25,26,48^, but also has been implicated in numerous anti-inflammatory effects^49,50,51,52^. Here, BioMapAI could identify a duality in its association with symptom severity. Similar biphasic relationships were observed for plasma metabolomics biomarkers, glucuronide and glutamine, in relation to pain (Figure 3C).

Distinct from other ‘omics features, KEGG genes exhibited sparse and dispersed contributions (Figure 3C, Supplemental Figure 4C). The vast feature matrix of KEGG models complicated the identification of a universal biomarker for any single symptom, as individuals possessed distinct symptom-specific KEGG biomarkers. For example, the gene FNR, an anaerobic regulatory protein transcription factor, was negatively associated with pain but appeared only in a small portion of patients, with the majority showing no significant impact (Figure 3C). This pattern was consistent for other KEGG biomarkers, which were sparsely associated with symptom severity (Supplemental Figures 4C).

Taken together, BioMapAI made associations between symptom-specific biomarkers and clinical phenotypes, which has been inaccessible to single models to date. Our models unveil a nuanced correlation between ‘omics features and disease symptomology, emphasizing ME/CFS’ complex etiology.

### Healthy Microbiome-Immune-Metabolome Networks are Dysbiotic in ME/CFS

BioMapAI elucidated that each ‘omics layer provided distinct insights into the disease symptoms and influenced host phenotypes in a dynamic and complex manner. To examine crosstalk between ‘omics layers, we modeled co-expression modules for each ‘omics using weighted gene co-expression network analysis (WGCNA), identifying seven microbial species, six microbial gene set, nine metabolome, and nine immune clusters (Methods, Supplemental Table 7). Observing significant associations of these modules with disease classification (microbial modules), age and gender (immune and metabolome modules) (Supplemental Figure 6A), we first established baseline networks of inter-‘omics interactions by calculating Spearman correlation coefficients (corrected, see Methods) among the module eigengenes of each omics cluster. An adjacency matrix was constructed using a cutoff of 0.3 to identify meaningful correlations, focusing on healthy individuals and incorporating clinical covariates such as age, weight, and gender (Figure 4A). We then examined how these correlations were altered in patient populations (Figure 4B, Supplemental Figure 6B-C).

**Figure 4:**
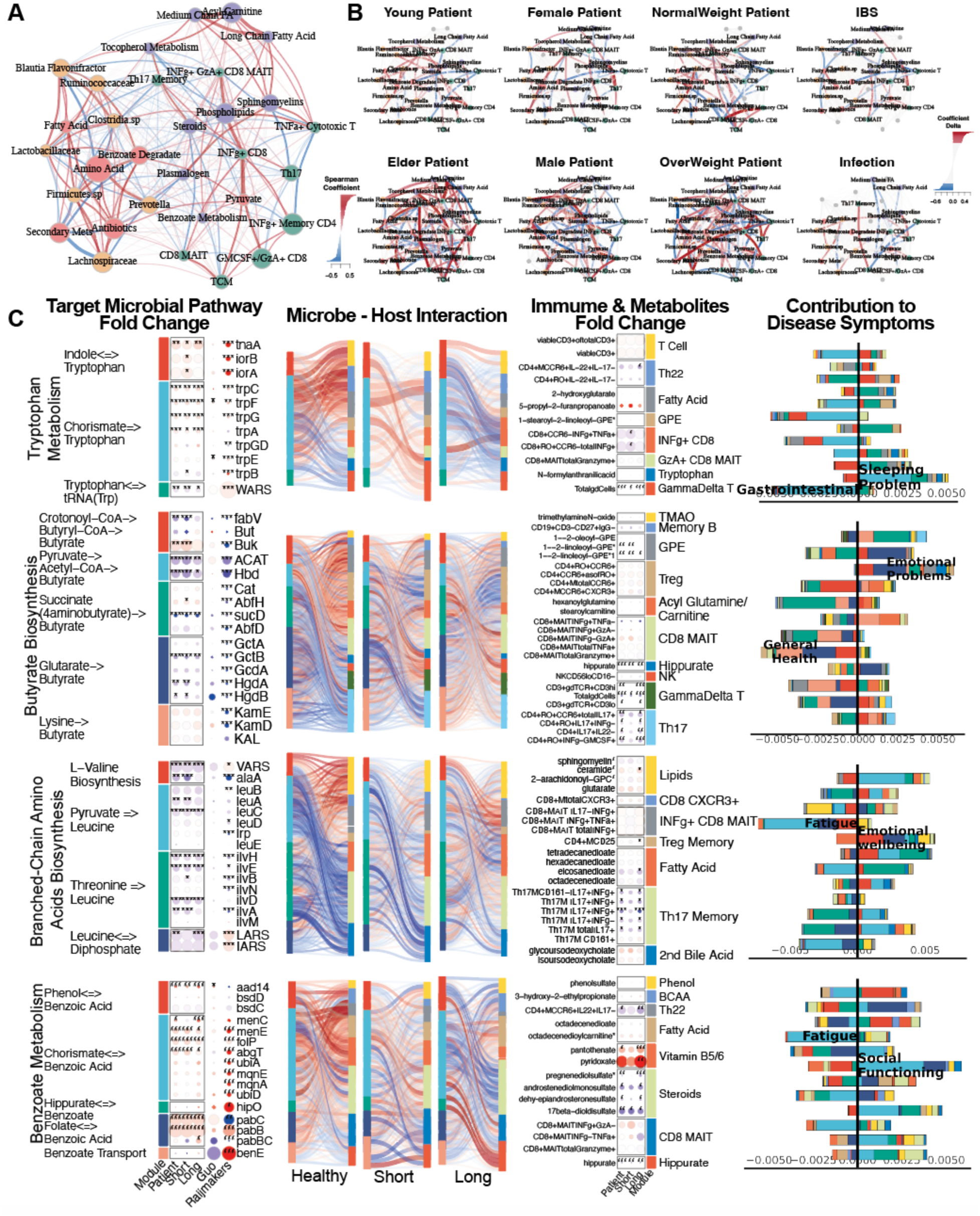
Microbiome-Immune-Metabolome Crosstalk is Dysbiotic in ME/CFS. A-B) Microbiome-Immune-Metabolome Network in A) Healthy and B) Patient Subgroups. A baseline network was established with 200+ healthy control samples (**A**), bifurcating into two segments: the gut microbiome (species in yellow, genetic modules in orange) and blood elements (immune modules in green, metabolome modules in purple). Nodes: modules; size: # of members; colors: ‘omics type; edges: interactions between modules, with Spearman coefficient (adjusted) represented by thickness, transparency, and color - positive (red) and negative (blue). Here, key microbial pathways (pyruvate, amino acid, and benzoate) interact with immune and metabolome modules in healthy individuals. Specifically, these correlations were disrupted in patient subgroups (**B**), as a function of gender, age (young <26 years old vs. older >50), BMI (normal <26 vs. overweight >26), and health status (individuals with IBS or infections). Correlations significantly shifted from healthy counterparts (Supplemental Figure 6C) are highlighted with colored nodes and edges indicating increased (red) or decreased (blue) interactions. **C) Targeted Microbial Pathways and Host Interactions.** Four microbial metabolic mechanisms (tryptophan, butyrate, BCAA, benzoate) were further analyzed to compare control, short and long-term ME/CFS patients, and external cohorts for validation (Guo^25^ and Raijmakers^26^).*1. Microbial Pathway Fold Change:* Key genes were grouped and annotated in subpathways. Circle size: fold change over control; color: increase (red) or decrease (blue), p-values (Patient vs Control, Wilcoxon, FDR adjusted) marked. *2. Microbiome-Host Interactions:* Sankey diagrams visualize interactions between microbial pathways and host immune cells/metabolites. Line thickness and transparency: Spearman coefficient (adjusted); color: red (positive), blue (negative). *3. Immune & Metabolites Fold Change:* Pathway-correlated immune cells and metabolites are grouped by category. *4. Contribution to Disease Symptoms:* Stacked bar plots show accumulated SHAP values (contributions to symptom severity) for each disease symptom (1-12, as in Supplemental Table 1). Colors: microbial subpathways and immune/metabolome categories match module color in fold change maps. X-axis: accumulated SHAP values (contributions) from negative to positive, with the most contributed symptoms highlighted. **P-values:** *p < 0.05, **p < 0.01, ***p < 0.001. **Abbreviations:** IBS, Irritable Bowel Syndrome; BMI, Body Mass Index; BCAA, Branched-Chain Amino Acids; MAIT, Mucosal-Associated Invariant T cell; SHAP, SHapley Additive exPlanations; GPE, Glycerophosphoethanolamine; INFγ, Interferon Gamma; CD, Cluster of Differentiation; Th, T helper cell; TMAO, Trimethylamine N-oxide; KEGG, Kyoto Encyclopedia of Genes and Genomes. **Supporting Materials:** Supplemental Table 7-8, Supplemental Figure 6.

Healthy control-derived host-microbiome interactions, such as the microbial pyruvate module associating with multiple immune modules, and connections between commensal gut microbes (*Prevotella*, *Clostridia* sp., *Ruminococcaceae*) with Th17 memory cells, plasma steroids, phospholipids, and tocopherol (vitamin E) (Figure 4A), were disrupted in ME/CFS patients.

Increased correlations between gut microbiome and mucosal/inflammatory immune modules, including CD8+ MAIT, and INFg+ CD4 memory cells, suggested an increased association with microbiome and inflammatory elements in ME/CFS (Supplemental Figure 6D). Young, female, and normal-weight patients shared those changes, while male patients showed different correlations between microbial and plasma metabolites. Elderly and overweight patients had more interaction abnormalities than other subgroups, with specific increases between *Blautia*, *Flavonifractor*, *Firmicutes* sp. linked with TNFα cytotoxic T cells and plasma plasmalogen, and decreased correlations between *Lachnospiraceae* sp. with Th17 cells (Figure 4B).

Further examining the pyruvate hub as well as several other key microbial modules whose networks were dysbiotic in patients, we mapped the correlations of their metabolic subpathways to plasma metabolites and immune cells and detailed the collective associations with host phenotypes (Figure 4C, Supplemental Table 8). We further validated these findings with two independent cohorts (Guo 2023^25^ and Raijmakers 2020^26^). For example, increased tryptophan metabolism, associated with gastrointestinal issues, lost its negative association with Th22 cells, and gained correlations with γδ T cells and the secretion of INFg and GzA from CD8 and CD8+ MAIT cells. Several networks associated with emotional dysregulation and fatigue – again underscoring the gut-brain axis^44^ – differed significantly in patients vs. controls, including decreased butyrate production - especially from the pyruvate^53^ and glutarate^54^ sub-pathways- and branched-chain amino acid (BCAA) biosynthesis, which had opposite correlations with Th17, Treg cells, and plasma lipids while having more correlations with inflammatory immune cells including γδ T and CD8+ MAIT cells in patients; and increased microbial benzoate, synthesized by *Clostridia* sp.^55,56^ then converted to hippurate in the liver^57,58^, showed a strong positive correlation with plasma hippurate in long-term ME/CFS patients, supporting enhanced pathway activity in later stages of the disease. These disrupted pathways also had modified associations with a variety of plasma metabolites—among them steroids, phenols, branched-chain amino acids, fatty acids, and vitamins B5 and B6. Notably, short-term ME/CFS patients presented a transitional profile, in which some health-associated networks were already dysbiotic but had not yet fully stabilized; these pathological connections became more firmly established in long-term ME/CFS.

Based on BioMapAI’s predictions and subsequent network analyses, we propose that some of the disease-specific changes in ME/CFS arise from disrupted associations between the gut microbiome, immune system, and metabolome (Figure 5). Reduced relative abundances of key microbes—such as *Faecalibacterium prausnitzii*—and corresponding disturbances in microbial metabolic pathways (e.g., butyrate, tryptophan, and BCAA production) correlated with pain and gastrointestinal abnormalities in ME/CFS. In healthy controls, these microbial metabolites are associated with activity of mucosal immune cells, including Th17, Th22, and Treg cells. In ME/CFS, however, these regulatory networks break down, with heightened pro-inflammatory responses mediated by γδ T cells and CD8 MAIT cells producing IFNγ and GzA, which in turn were associated with subjective health perception and social functioning.

**Figure 5:**
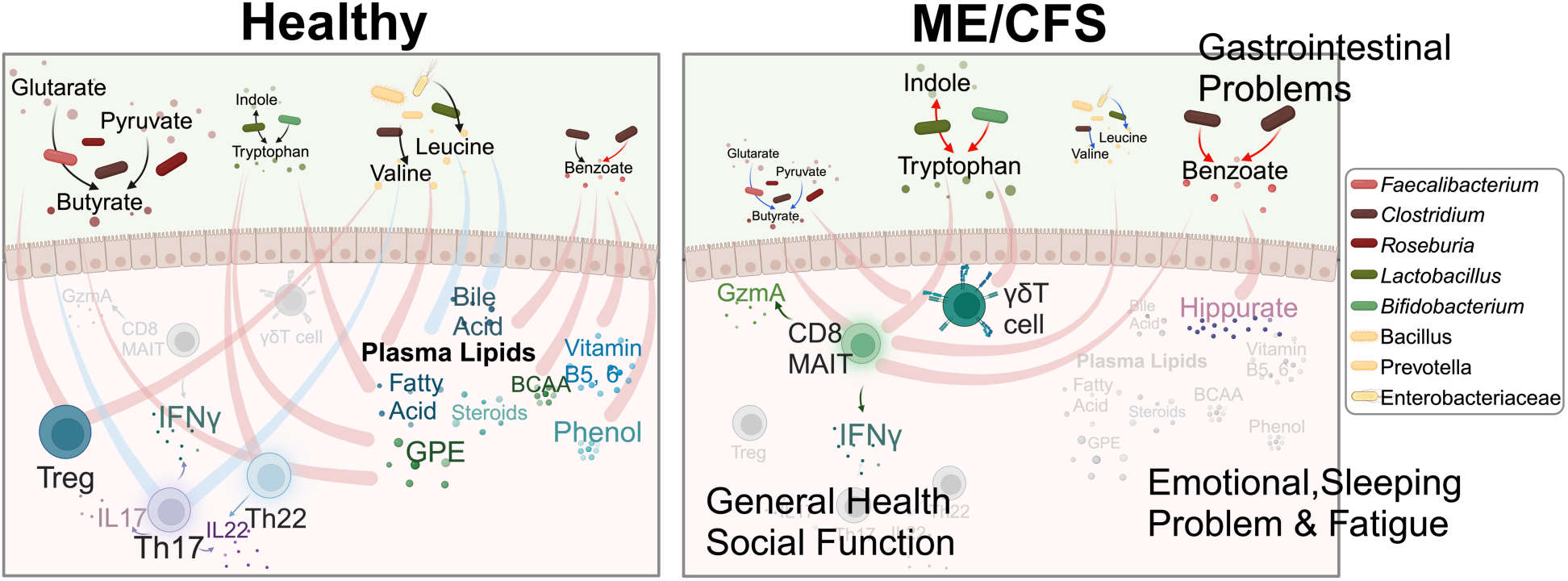
Overview of Dysbiotic Host-Microbiome Interactions in ME/CFS. This conceptual diagram visualizes the host-microbiome interactions in healthy conditions (left) and its disruption and transition into the disease state in ME/CFS (right). The base icons of the figure remain consistent, while gradients and changes in color and size visually represent the progression of the disease. Process of production and processing is represented by lines with arrows, where the color indicates an increase (red) or decrease (blue) in the pathway in disease; lines without arrows indicate correlations, with red representing positive and blue representing negative correlations. In healthy conditions, microbial metabolites support immune regulation, maintaining mucosal integrity and healthy inflammatory responses by positively regulating Treg and Th22 cell activity, and controlling Th17 activities, including the secretion of IL17 (purple cells), IL22 (blue), and IFNγ. These microbial metabolites also maintain many positive interactions with plasma metabolites like lipids, bile acids, vitamins, and phenols. In ME/CFS, there is a significant decrease in beneficial microbes and a disruption in metabolic pathways, marked by a decrease in the butyrate (brown-red dots) and BCAA (yellow) pathways and an increase in tryptophan (green) and benzoate (red) pathways. These changes are linked to gastrointestinal issues. In ME/CFS, the regulatory capacity of the immune system diminishes, leading to the loss of health-associated interactions with Th17, Th22, and Treg cells, and an increase in inflammatory immune activity. Pathogenic immune cells, including CD8 MAIT and γδT cells, show increased activity, along with the secretion of inflammatory cytokines such as IFNγ and GzmA, contributing to worsened general health and social functioning. Healthy interactions between gut microbial metabolites and plasma metabolites weaken or even reverse in the disease state. A notable strong connection increased in ME/CFS is benzoate transformation to hippurate, associated with emotional disturbances, sleep issues, and fatigue. **Abbreviations:** IFNγ, Interferon gamma; Th17, T helper 17 cells; Th22, T helper 22 cells; Treg, Regulatory T cells; GzmA, Granzyme A; MAIT, Mucosa-Associated Invariant T cells; γδT, Gamma delta T cells; BCAA, Branched-Chain Amino Acids; GPE, Glycerophosphoethanolamine.

Additional health-associated interactions between microbial benzoate metabolism and various plasma metabolites (e.g., lipids, glycerophosphoethanolamine, fatty acids, and bile acids) we hypothesized are also diminished or reversed in ME/CFS. This breakdown in host–microbiome metabolic networks correlates with more severe fatigue, emotional disturbances, and sleep problems, aligning with emerging evidence that microbially derived metabolites may affect the gut–brain axis^59,60,61^.

## Discussion

Democratization of AI technologies and large-scale multi-‘omics has the promise of revolutionizing precision medicine^62,63,64,65^. This study generated among the most extensive paired multi-’omics dataset for ME/CFS to date^66,25,26,27,28,67,68,69^, bringing new technical and biological insights. Technically, BioMapAI marks the first supervised deep learning model trained to accommodate these complex, multi-system ME/CFS symptoms. The rationale behind BioMapAI is that understanding long-term, post-infection syndromes like ME/CFS is not necessarily solved by pinpointing an exact diagnosis or tracing disease origins^70,2^,^71^, but rather by addressing the chronic, multifaceted symptoms that significantly impacts patients’ quality of life^72,73^. Biologically, our study introduces a highly nuanced approach to link physiological changes in gut microbiome, plasma metabolome, and immune status, with host symptoms, moving beyond the initial causes of the disease^74,75^. Importantly, we validated key biomarkers in external cohorts^25,26,27,28^, despite significant demographic and methodological differences between the studies.

This study represents a substantial technical and biological advance over our previous work and other investigations of ME/CFS to date. First, we developed BioMapAI, a supervised deep neural network architecture that accommodates the full complexity of our multi-omics datasets— encompassing gut microbiome, plasma metabolome, immune profiling, blood labs, and extensive clinical surveys—beyond what traditional ML models can handle. By jointly modeling these diverse data types, BioMapAI explains the phenotypic heterogeneity of ME/CFS more effectively than single-outcome methods and simultaneously identifies symptom-specific biomarkers. Furthermore, our dataset’s unprecedented size, in both participant numbers and the depth of datatypes, allowed us to build a robust AI model validated on both held-out data and external cohorts. As a sanity check, we confirmed key biomarkers—such as altered *Faecalibacterium prausnitzii* and butyrate producers (reported by Guo et al.) as well as sphingolipid pathway changes (described by Raijmakers et al., Germain et al., and Che et al.)— using independent datasets, which other studies have not performed. Nonetheless, a caveat of our model is that from a clinical perspective, simply distinguishing ME/CFS from healthy controls may be less challenging than differentiating ME/CFS from other conditions with overlapping symptoms, such as fibromyalgia. To establish whether our pre-trained model (“DeepMECFS”) can discriminate among multiple chronic diseases, similar datasets with other diseases and comparative models are needed in future work. Second, we added a new, detailed blood immune-profiling dataset, which provided the most biologically explanatory features for both disease classification and symptom severity.

Leveraging these data, we were able to construct new microbiome–metabolome–immune networks in both health and ME/CFS—an advance over earlier investigations that generally focused on only one ‘omics layer (e.g., stool microbiome in Guo et al.; plasma metabolomics in Germain et al. and Che et al.). While Raijmakers et al. examined 92 inflammatory circulating markers, plasma metabolites, and gut microbiome in a smaller study (n=50 ME/CFS, n=72 healthy control for metagenomics, and n=22 for metabolomics), their analyses were relatively limited in that they used ML models to differentiate ME/CFS from controls and only examined fatigue as a clinical variate, not adjusting for other clinical variables that could affect ‘omics associations such as age, gender, or BMI. Moreover, their approach only assessed pairwise associations among data types. In contrast, our multi-‘omics strategy explicitly accounts for demographic and clinical covariates like age, gender and BMI, revealing that these factors can markedly reshape immune–microbiome–metabolome interaction networks, just as comorbid conditions such as obesity or advanced age can further individualize disease phenotypes.

Taken together, our dataset uncovers an array of correlations that while not explaining causality or confirming mechanism, can further our understanding of ME/CFS in several ways. First, our analyses underscore the importance of considering clinical symptom heterogeneity and cohort-level covariates because interactions among the microbiome, metabolome, and immune system vary substantially depending on these factors. Although it has long been assumed that confounders play a major role, previous studies have seldom controlled for them in a comprehensive manner, potentially explaining some of the inconsistencies reported in single-‘omics analyses. Second, while our findings are correlative rather than causal, they generate numerous hypotheses about both specific and more extensive pathways that may be disrupted in ME/CFS. For example, our previous analysis, and work by Guo et al., suggest that diminished butyrate-producing microbes in ME/CFS lower the availability of short-chain fatty acids (SCFAs) in the stool (Guo) and plasma (Xiong). Here, we refine that hypothesis by pinpointing potential immunological or metabolic mediators of this change. In healthy controls, multiple butyrate biosynthesis routes are inversely associated with Th17 cells, whereas the glutarate→butyrate pathway aligns with Tregs. These patterns become largely reversed in long-term disease, with succinate→butyrate showing new negative correlations to Tregs and positive links with CD8+ MAIT cells. ME/CFS also substantially alters metabolite associations with Th17 cells.

On the metabolomic side, there is currently no direct biochemical link reported between glutarate→butyrate and glycerophosphoethanolamine (GPE)—though in healthy controls, they exhibited a strong positive correlation which was altered in ME/CFS. One can then hypothesize an indirect link with phospholipid metabolism and its effect on neurotransmission.

In addition to refining established hypotheses, our results propose new links among tryptophan metabolism, branched-chain amino acids (BCAAs), and benzoate metabolism in shaping immune function and symptomatology in ME/CFS. Although no direct biochemical connection between tryptophan metabolism and 2-hydroxyglutarate is currently known, both pathways likely influence immune regulation and metabolic reprogramming, indicating a more complex regulatory landscape. In healthy controls, tryptophan metabolism is closely tied to various T cell subsets, including Th22 cells, whereas these relationships are disrupted in ME/CFS. Furthermore, we observed significant alterations in benzoate metabolism modules and their associations with plasma steroids, hippurate, and fatty acids. These pathways, linked to both steroid biosynthesis and neurotransmitter production (e.g., serotonin, cortisol), highlight a potential gut–brain axis component in ME/CFS pathophysiology.

While some of these findings may seem granular or only indirectly testable—such as potential sex differences in the interaction network—our detailed, multi-‘omics perspective is valuable for unraveling the disease’s heterogeneity. As experimental models attempt to validate these hypotheses, one must keep in mind that many interactions may be context- or model-specific rather than universally turned on or off in disease states. This context dependency underscores the need for nuanced, carefully controlled mechanistic studies that incorporate patient heterogeneity and environmental factors when investigating ME/CFS.

Additional limitations of our study include that that our study population was comprised more females and older individuals, primarily Caucasian, though this is consistent with the epidemiology of ME/CFS^18,76,77^, and was from a single geographic location (Bateman Horne Center). This may limit our findings to certain populations. In addition, previous RNA sequencing studies have suggested mitochondrial dysfunction and altered energy metabolism in ME/CFS^78,79,80,81,82^; thus, incorporating host PBMC RNA or ATAC sequencing in future research could provide deeper insights into regulatory changes. The typical decades-long disease progression of ME/CFS makes it challenging for our four-year longitudinal design to capture stable temporal signals - although separating our short-term (<4 years) and long-term (>10 years) provided valuable insights – ideally, tracking the same patients over a longer period would likely yield more accurate trends^83,84^. Long disease history also increases the likelihood of exposure to various diets and medications^85^, which could influence biomarker identification, particularly in metabolomics. Finally, model-wise, BioMapAI was trained on < 500 samples with tenfold cross-validation, which is relatively small given the complexity of the outcome matrix; expanding the training dataset and incorporating more independent validation sets could potentially enhance its performance and generalizability^86,87^. Currently, the model treated all 12 studied symptoms with equal importance due to the unclear symptom prioritization in ME/CFS^88^. We computed modules to assign different weights to symptoms to enhance diagnostic accuracy. While this approach was not particularly effective for ME/CFS, it may be more promising for diseases with more clearly defined symptom hierarchies^89,90^. In such cases, adjusting the weights of symptoms in the model’s final layer could improve performance and help pinpoint which symptoms more strongly contributing.

Although our findings are still preliminary for direct therapeutic application, the nuanced insights and deconstructed approach described here offer numerous hypotheses for dysbiotic microbiome–metabolome–immune connections in ME/CFS. We hope that the unprecedented systems-level resolution of our dataset, algorithm, and analyses will contribute to filling out heretofore unknown links between these factors thus explaining some of the disease heterogeneity in this important disease.

## Methods

### Study Design

This was 4-year prospective study. All participants had a physical examination at the baseline visit that included evaluation of vital signs, BMI, orthostatic vital signs, skin, lymphatic system, HEENT, pulmonary, cardiac, abdomen, musculoskeletal, nervous system and fibromyalgia (FM) tender points. We enrolled a total of 153 ME/CFS patients (of which 75 had been diagnosed with ME/CFS <4 years before recruitment and 78 had been diagnosed with ME/CFS >10 years before recruitment) and 96 healthy controls. Among them, 110 patients and 58 healthy controls were followed one year after the recruitment as timepoint 2; 81 patients and 13 healthy controls were followed two years after the recruitment as timepoint 3; and 4 patients were followed four years after the recruitment as timepoint 4. Subject characteristics are shown in Supplemental Table 1 and Supplemental Figure 1A.

Medical history and concomitant medications were documented. Blood samples were obtained prior to orthostatic and cognitive testing. The 10-minute NASA Lean Test and cognitive testing were conducted after the physical examination and blood draw^91^. Cognitive efficiency was tested with the DANA Brain Vital, measuring three reaction time and information processing measurements^92^. The orthostatic challenge was assessed with the 10-minute NASA Lean Test (NLT). Participants rested supine for 10 minutes, and baseline blood pressure (BP) and heart rate (HR) were measured twice during the last 2 minutes of rest^93^.

Participants were provided with an at-home stool collection kit at the end of each in-person visit. The following questionnaires were completed at baseline: DePaul Symptom Questionnaire (DSQ), Post-Exertional Fatigue Questionnaire, RAND-36, Fibromyalgia Impact Questionnaire-R, ACR 2010 Fibromyalgia Criteria Symptom Questionnaire, Pittsburgh Sleep Quality Index (PSQI), Stanford Brief Activity Survey, Orthostatic Intolerance Daily Activity Scale, Orthostatic Intolerance Symptom Assessment, Brief Wellness Survey, Hours of Upright Activity (HUA), medical history and family history. All but medical history and family history were administered again when participants came for their annual visit.

Approval was received before enrolling any subjects in the study (The Jackson Laboratory Institutional Review Board, 17-JGM-13). All participants were educated about the study prior to enrollment and signed all appropriate informed consent documents. Research staff followed Good Clinical Practices (GCP) guidelines to ensure subject safety and privacy.

### ME/CFS Cohort

Beginning in January 2018, we enrolled ME/CFS patients who had been sick for <4 years or sick for >10 years. No ME/CFS patients with duration ≥4 years and ≤10 years were enrolled in order to have clear distinctions between short and long duration of illness with ME/CFS. All participants were 18 to 65 years old at the time of enrollment. ME/CFS diagnosis according to the Institute of Medicine clinical diagnostic criteria and disease duration of <4 years were confirmed during clinical differential diagnosis and thorough medical work up^94^. Additional inclusion criteria required, 1) a substantial reduction or impairment in the ability to engage in pre-illness levels of occupational, educational, social, or personal activities that persists for more than 6 months and less than 4 years and is accompanied by fatigue, which is often profound, is of new or definite onset (not lifelong), is not the result of ongoing excessive exertion, and is not substantially alleviated by rest, and 2) post-exertional malaise. Exclusionary criteria for the <4 year ME/CFS cohort were, 1) morbid obesity BMI>40, 2) other active and untreated disease processes that explain most of the major symptoms of fatigue, sleep disturbance, pain, and cognitive dysfunction, 3) untreated primary sleep disorders, 4) rheumatological disorders, 5) immune disorders, 6) neurological disorders, 7) infectious diseases, 8) psychiatric disorders that alter perception of reality or ability to communicate clearly or impair physical health and function, 9) laboratory testing or imaging are available that support an alternate exclusionary diagnosis, and 10) treatment with short-term (less than 2 weeks) antiviral or antibiotic medication within the past 30 days.

For the >10 year ME/CFS cohort, disease duration of >10 year and clinical criteria was confirmed to meet the Institute of Medicine criteria for ME/CFS during clinical evaluation and medical history review^94^. Other than disease duration, inclusion and exclusion criteria were the same as for <4 year ME/CFS cohort.

### Healthy Control Cohort

Healthy control participants were also between 18 to 65 years of age and in general good health. Enrollment began in 2018 and subjects were selected to match the <4 year ME/CFS cohort by age (within 5 years), race, and sex (∼2:1 female to male ratio). Exclusion criteria for healthy controls included, 1) a diagnosis or history of ME/CFS, 2) morbid obesity BMI>40, 3) treatment with short-term (less than 2 weeks) antiviral or antibiotic medication within the past 30 days or 4) treatment long-term (longer than 2 weeks) antiviral medication or immunomodulatory medications within the past 6 months.

### Clinical Metadata and Scores

Clinical symptoms and baseline health status were assessed on the day of physical examination and biological sample collection for both case and control subjects. For each participant, we collected demographic information (including age, gender, diet, race, BMI, family, work, and education), medical histories, clinical tests and questionnaires. From questionnaires and test as described above, we summarized 12 clinical scores to cover major symptoms of ME/CFS: Scores 1-8 were derived from the RAND36, following standardized rules ^95^ and summarized into eight categories: Physical Functioning (also referred to as Daily Activity in the main contents), Role Limitations due to Physical Health (Physical Limitations), Role Limitations due to Emotional Problems (Emotional Problems), Energy/Fatigue, Emotional Wellbeing (Mental Health), Social Functioning (Social Activity), Pain, and General Health (Health Perception). Cognitive Efficiency was summarized from the DANA Brain Vital test, Orthostatic Intolerance from the NLT test, Sleeping Problem Score from the Pittsburgh Sleep Quality Index (PSQI) questionnaire, and Gastrointestinal Problems Score from the Gastrointestinal Symptom Rating Scale (GSRS) questionnaire. Each score was transformed into a 0–1 scale to facilitate combination and comparison, where a score of 1 indicates maximum disability or severity and a score of 0 indicates no disability or disturbance.

### Plasma Sample collection and Preparation

Healthy and patient blood samples were obtained from Bateman Horne Center, Salt Lake City, UT and approved by JAX IRB. One 4 mL lavender top tube (K2EDTA) was collected, and tube slowly inverted 8-10 times immediately after collection. Blood was centrifuged within 30 minutes of collection at 1000 x g with low brake for 10 minutes. 250 uL of plasma was transferred into three 1 mL cryovial tubes, and tubes were frozen upright at −80°C. Frozen plasma samples were batch shipped overnight on dry ice to The Jackson Laboratory, Farmington, CT, and stored at −80°C. Heparinized blood samples were shipped overnight at room temperature. Peripheral blood mononuclear cells (PBMC) were isolated using Ficoll-paque plus (GE Healthcare) and cryopreserved in liquid nitrogen.

### Plasma untargeted metabolome by UPLC-MS/MS

Plasma samples were sent to Metabolon platform and processed by Ultrahigh Performance Liquid Chromatography-Tandem Mass Spectroscopy (UPLC-MS/MS) following the CFS cohort pipeline. In brief, samples were prepared using the automated MicroLab STAR® system from Hamilton Company. The extract was divided into five fractions: two for analysis by two separate reverse phases (RP)/UPLC-MS/MS methods with positive ion mode electrospray ionization (ESI), one for analysis by RP/UPLC-MS/MS with negative ion mode ESI, one for analysis by HILIC/UPLC-MS/MS with negative ion mode ESI, and one sample was reserved for backup. QA/QC were analyzed with several types of controls were analyzed including a pooled matrix sample generated by taking a small volume of each experimental sample (or alternatively, use of a pool of well-characterized human plasma), extracted water samples, and a cocktail of QC standards that were carefully chosen not to interfere with the measurement of endogenous compounds were spiked into every analyzed sample, allowed instrument performance monitoring, and aided chromatographic alignment.

Compounds were identified by comparison to Metabolon library entries of purified standards or recurrent unknown entities. The output raw data included the annotations and the value of peaks quantified using area-under-the-curve for metabolites.

### Immune Profiling: Flow Cytometry Analysis

Frozen PBMC aliquots were thawed, counted and divided into two parts, one part for day 0 surface staining, and the other part cultured in complete RPMI 1640 medium (RPMI plus 10% Fetal Bovine Serum (FBS, Atlanta Biologicals) and 1% penicillin/streptomycin (Corning Cellgro) supplemented with IL-2+IL15 (20ng/ml) for Treg subsets day 1 surface and transcription factors staining after culture with IL-7 (20ng/ml) for day 1 and day 6 intracellular cytokine staining, and a combination of cytokines (20ng/ml IL-12, 20ng/ml IL-15, and 40ng/ml IL-18) for day 1 intracellular cytokine staining (IL-12 from R&D, IL-7 and IL-15 from Biolegend). Surface staining was performed in staining buffer containing PBS + 2% FBS for 30 minutes at 4°C. When staining for chemokine receptors the incubation was done at room temperature. Antibodies used in the surface staining are 2B4, CD1c, CD14, CD16, CD19, CD25, CD27, CD31, CD3, CD303, CD38, CD4, CD45RO, CD56, CD8, CD95, CD161, CCR4, CCR6, CCR7, CX3CR1, CXCR3, CXCR5, γδ TCR bio, HLA-DR, IgG, IgM, LAG3, PD-1, TIM3, Va7.2, Va24Ja18 all were obtained from Biolegend.

For intracellular cytokine staining, cells were stimulated with PMA (40ng/ml for overnight cultured cells and 20ng/ml for 6 days cultured cells) and Ionomycin (500ng/ml) (both from Sigma-Aldrich) in the presence of GolgiStop (BD Biosciences) for 4 hours at 37°C. For cytokine secretion after stimulation with IL-12+IL-15+IL-18, GolgiStop was added to the culture on day 1 for 4 hours. For intracellular cytokine and transcription factor staining, PMA+Ionomycin stimulated cells of unstimulated cells were collected, stained with surface markers including CD3, CD4, CD8, CD161, PD1, 2B4, Vα7.2, CD45RO, CCR6, and CD27 followed by one wash with PBS (Phosphate buffer Saline) and staining with fixable viability dye (eBioscience). After surface staining, cells were fixed and permeabilized using fixation/permeabilization buffers (eBioscience) according to the manufacturer’s instruction. Permeabilized cells were then stained for intracellular FOXP3, Helios, IL-4, IFNγ, TNFα, IL-17A, IL-22, Granzyme A, GM-CSF, and Perforin from Biolegend. Flow cytometry analysis was performed on Cytek Aurora (Cytek Biosciences) and analyzed using FlowJo (Tree Star).

### Fecal Sample Collection and DNA Extraction

Stool was self-collected at home by volunteers using a BioCollector fecal collection kit (The BioCollective, Denver, CO) according to manufacturer instructions for preservation for sequencing prior to sending the sample in a provided Styrofoam container with a cold pack. Upon receipt, stool and OMNIgene samples were immediately aliquoted and frozen at –80°C for storage. Prior to aliquoting, OMNIgene stool samples were homogenized by vortexing (using the metal bead inside the OMNIgene tube), then divided into 2 microfuge tubes, one with 100µL aliquot and one with 1mL. DNA was extracted using the Qiagen (Germantown, MD, USA) QIAamp 96 DNA QIAcube HT Kit with the following modifications: enzymatic digestion with 50μg of lysozyme (Sigma, St. Louis, MO, USA) and 5U each of lysostaphin and mutanolysin (Sigma) for 30 min at 37 °C followed by bead-beating with 50 μg 0.1 mm of zirconium beads for 6 min on the Tissuelyzer II (Qiagen) prior to loading onto the Qiacube HT. DNA concentration was measured using the Qubit high sensitivity dsDNA kit (Invitrogen, Carlsbad, CA, USA).

### Metagenomic Shotgun Sequencing

Approximately 50µL of thawed OMNIgene preserved stool sample was added to a microfuge tube containing 350 µL Tissue and Cell lysis buffer and 100 µg 0.1 mm zirconia beads. Metagenomic DNA was extracted using the QiaAmp 96 DNA QiaCube HT kit (Qiagen, 5331) with the following modifications: each sample was digested with 5µL of Lysozyme (10 mg/mL, Sigma-Aldrich, L6876), 1µL Lysostaphin (5000U/mL, Sigma-Aldrich, L9043) and 1µL oh Mutanolysin (5000U/mL, Sigma-Aldrich, M9901) were added to each sample to digest at 37°C for 30 minutes prior to the bead-beating in the in the TissueLyser II (Qiagen) for 2 x 3 minutes at 30 Hz. Each sample was centrifuged for 1 minute at 15000 x g prior to loading 200µl into an S-block (Qiagen, 19585) Negative (environmental) controls and positive (in-house mock community of 26 unique species) controls were extracted and sequenced with each extraction and library preparation batch to ensure sample integrity. Pooled libraries were sequenced over 13 sequencing runs using both HiSeq (N=87) and NovaSeq (N=392) platforms. To address potential biases arising from varying read depths, all samples were down-sampled, using seqtk^96^ (v1.3-r106), to 5 million reads. This threshold corresponds to the 95th percentile of the read count distribution across the dataset.

Sequencing adapters and low-quality bases were removed from the metagenomic reads using scythe (v0.994) and sickle (v1.33), respectively, with default parameters. Host reads were removed by mapping all sequencing reads to the hg19 human reference genome using Bowtie2 (v2.3.1), under ‘very-sensitive’ mode. Unmapped reads (i.e., microbial reads) were used to estimate the relative abundance profiles of the microbial species in the samples using MetaPhlAn4.

### Taxonomic Profiling (Specie Abundance) and KEGG Gene Profiling

Taxonomic compositions were profiled using Metaphlan4.0^97^ and the species whose average relative abundance > 1e-4 were kept for further analysis, giving 384 species. The gene profiling was computed with USEARCH^98^ (v8.0.15) (with parameters: evalue 1e-9, accel 0.5, top_hits_only) to KEGG Orthology (KO) database v54, giving a total of 9452 annotated KEGG genes. The reads count profile was normalized by DeSeq2^99^ in R. Genes with a prevalence of over 20% were selected for downstream analysis.

### Confounder Analysis

Confounder analysis was done by R package MaAsLin2^100^. We considered demographic features (including age, gender, BMI, ethnicity, and race), diet records, medications (antivirals, antifungals, antibiotics, and probiotics), and self-reported IBS scores as potential confounders. The analysis followed the model formula:

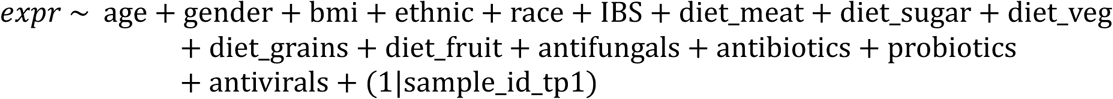

where *expr* refers to the ‘omics matrix. For each feature in the ‘omics data, we ran this generalized linear model to identify multivariable associations between each ‘omics feature and each metadata feature. Identified confounders were handled differently based on the type of data. For species and KEGG genes, any feature with a significant statistical association with any metadata feature was removed from all subsequent analyses, resulting in the removal of 21 species and 946 microbial genes. For immune profiling and plasma metabolomics, to remove the effects of identified confounders, each feature was adjusted by retaining the residuals^97^, i.e., the part of the outcome not explained by the confounding factors, from a general linear model:

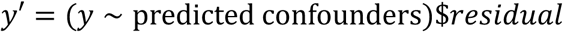

Additionally, for network and patient subset analysis (Methods), age, gender, BMI, and IBS were not included as confounders since we analyzed different age groups, gender groups, weight groups, and IBS groups separately. However, other identified confounders were still considered in the residual models.

### BioMapAI

The rationale behind BioMapAI is we believe that ME/CFS is characterized by significant heterogeneity and individual variability, making traditional approaches—such as classifying patients versus controls and reporting single-disease biomarkers—insufficient to us. This motivated us to develop a sophisticated model that directly integrates rich biological multi-omics data with clinical phenotypes. The primary learning goal of BioMapAI is to connect high-dimensional biology data, *X* to mixed-type output matrix, *Y*. Unlike traditional ML or DL classifiers that typically predict a single outcome, *y*, BioMapAI is designed to learn multiple objects, *Y* = [*y*_1_, *y*_2_, …, *y*_*n*_], simultaneously within a single model. This approach allows for the simultaneous prediction of diverse clinical outcomes - including binary, categorical, continuous variables - with ‘omics profiles, thus address disease heterogeneity by tailoring each patient’s specific symptomology. The uniqueness of BioMapAI is it is the first supervised deep learning model that integrates omics directly with clinical phenotypes in ME/CFS. This design enables simultaneous identification of symptom-specific and disease-general biomarkers, accounting for ME/CFS’s phenotypic heterogeneity.

#### 1. BioMapAI Structure

BioMapAI is a fully connected deep neural network framework comprising an input layer *X*, a normalization layer, three sequential hidden layers, *Z*^1^, *Z*^2^, *Z*^3^,and one output layer *Y*.

1. **Input layer (***X*) takes high-dimensional ‘omics data, such as gene expression, species abundance, metabolome matrix, or any customized matrix like immune profiling and blood labs.
2. **Normalization Layer** standardizes the input features to have zero mean and unit variance, defined as

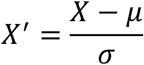

where μ is the mean and σ is the standard deviation of the input features.
3. **Feature Learning Module** is the core of BioMapAI, responsible for extracting and learning important patterns from input data. Each fully connected layer (hidden layer 1-3) is designed to capture complex interactions between features. **Hidden Layer 1 (**Z^1^) and **Hidden Layer 2 (**Z^2^) contain 64 and 32 nodes, respectively, both with ReLU activation and a 50% dropout rate, defined as:

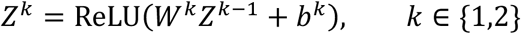

**Hidden Layer 3 (**Z^3^) has *n* parallel sub-layers for each object, *y*_*i*_ in *Y*. Every sub-layer, *Z*^3^, contains 8 nodes, represented as:

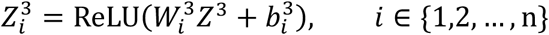

All hidden layers used ReLU activation functions, defined as:

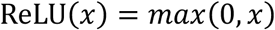
4. **Outcome Prediction Module** is responsible for the final prediction of the objects. **The output layer (***Y*) has *n* nodes, each representing a different object:

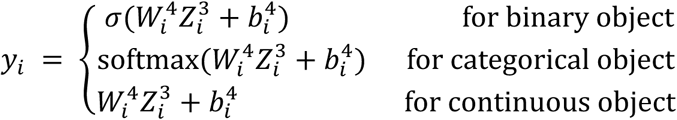

The loss functions are dynamically assigned based on the type of each object:

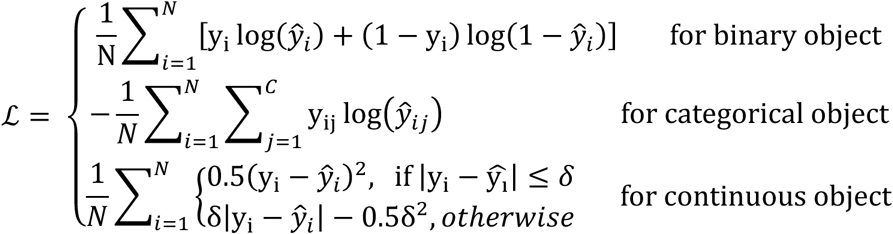

During training, the weights are adjusted using the Adam optimizer. The learning rate was set to 0.01, and weights were initialized using the He normal initializer. L2 regularizations were applied to prevent overfitting.
5. **Optional Binary Classification Layer** (not used for parameter training). An additional binary classification layer is attached to the output layer *Y* to evaluate the model’s performance in binary classification tasks. This layer is not used for training BioMapAI but serves as an auxiliary component to assess the accuracy of predicting binary outcomes, for example, disease vs. control. This ScoreLayer takes the predicted scores from the output layer and performs binary classification:

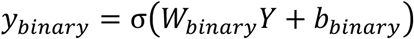

The initial weights of the 12 scores are derived from the original clinical data, and the weights are adjusted based on the accuracy of BioMapAI’s predictions:

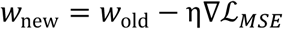

where ∇ℒ_*MSE*_ refers to the mean squared error (MSE) between the predicted *y*’ and true *y*, then adjusts the weights to optimize the accuracy of the binary classification.

#### 2. Training and Evaluation of BioMapAI for ME/CFS – BioMapAI::DeepMECFS

BioMapAI is a framework designed to connect high-dimensional, sparse biological ‘omics matrix *X* to multi-output *Y*. While BioMapAI is not tailored to a specific disease, it is versatile and applicable to a broad range of biomedical topics. In this study, we trained and validated BioMapAI using our ME/CFS datasets. The trained models are available on GitHub, nicknamed DeepMECFS, for the benefit of the ME/CFS research community.

1. **Dataset Pre-Processing Module: Handling Sample Imbalance.** To ensure uniform learning for each output *y*, it is crucial to address sample imbalance before fitting the framework. We recommend using customized sample imbalance handling methods, such as Synthetic Minority Over-sampling Technique (SMOTE)^101^, Adaptive Synthetic (ADASYN)^102^, or Random Under-Sampling (RUS)^103^. In our ME/CFS dataset, there is a significant imbalance, with the patient data being twice the size of the control data. To effectively manage this class imbalance, we employed RUS as a random sampling method for the majority class. Specifically, we randomly sampled the majority class 100 times. For each iteration *i*, a different random subset *S*_i_^*majority*^ was used. This subset *S*^*majority*^ of the majority class was combined with the entire minority class *S*_i_^*minority*^. For each iteration *i*:

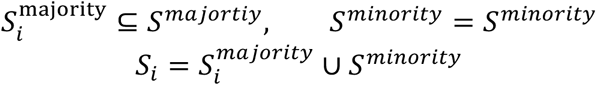

where the combined dataset *S*_*i*_ was used for training at each iteration. This approach allows the model to generalize better and avoid biases towards the majority class, improving overall performance and robustness.
2. **Model Training, Cross-Validation and Held-out Validation.** DeepMECFS is the name of the trained BioMapAI model with ME/CFS datasets. We trained on five preprocessed ‘omics datasets, including species abundances (Feature N=118, Sample N=474) and KEGG gene abundances (Feature N=3959, Sample N=474) from the microbiome, plasma metabolome (Feature N=730, Sample N=407), immune profiling (Feature N=311, Sample N=481), and blood measurements (Feature N=48, Sample N=495). Additionally, an integrated ‘omics profile was created by merging the most predictive features from each ‘omics model related to each clinical score (SHAP Methods), forming a comprehensive matrix of 154 features, comprising 50 immune features, 32 species, 30 KEGG genes, and 42 plasma metabolites. To evaluate the performance of BioMapAI, we employed a robust 10-fold cross-validation alongside a held-out validation approach. Specifically, 10% of the data was excluded from the cross-validation process to serve as an independent validation set. This allowed us to assess both the model’s performance during cross-validation and its generalizability on unseen data. Training was conducted over 500 epochs with a batch size of 64 and a learning rate of 0.0005, optimized through grid search. The Adam optimizer was used to adjust the weights during training, chosen for its ability to handle sparse gradients on noisy data. The initial learning rate was set to 0.0005, with beta1 set to 0.9, beta2 set to 0.999, and epsilon set to 1e-7 to ensure numerical stability. Dropout layers with a 50% dropout rate were used after each hidden layer to prevent overfitting, and L2 regularization (λ = 0.008) was applied to the kernel weights, defined as:

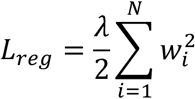
3. **Model Evaluation.** To evaluate the performance of the models, we employed several metrics tailored to both regression and classification tasks. The Mean Squared Error (MSE) was used to evaluate the performance of the reconstruction of each object. For each *y*_*i*_, MSE was calculated as:

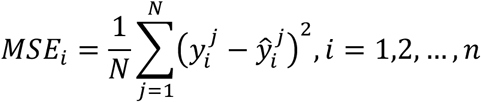

where *y*^j^_i_ is the actual values, Ŷ_i_^j^ is the predicted values, and *N*_*i*_ is the number of samples, *n*_*i*_ is the number of objects. For binary classification tasks (ME/CFS vs control), we utilized multiple metrics including accuracy, precision, recall, and F1 score to enable a comprehensive evaluation of the model’s performance. To benchmark the performance of BioMapAI, we compared its binary classification performance with four traditional machine learning models and one deep neural network (DNN) model. The traditional machine learning models included: 1) Logistic Regression (**LR**) (C=0.5, saga solver with Elastic Net regularization); 1) Generalized linear modeling with elastic net regularization (**Glmnet**) (grid search for best alpha/lambda, tuneLength = 10) - R glmnet, caret; 2) Glmnet with interaction terms (**Glmnet-int**) - R glmnet, caret; 3) Support Vector Machine (**SVM**) with an RBF kernel (C=2) - sklearn.svm.SVC; and 4) Gradient Boosting Decision Trees (**GBDT**) (learning rate = 0.05, maximum depth = 5, estimators = 1000) - sklearn.ensemble.GradientBoostingClassifier. **DNN** model employed the same hyperparameters as BioMapAI, except it did not include the parallel sub-layer, *Z*_3_, thus it only performed binary classification instead of multi-output predictions. The comparison between BioMapAI and DNN aims to assess the specific contribution of the spread-out layer, designed for discerning object-specific patterns, in binary prediction. Evaluation metrics are detailed in Supplemental Table 3.
4. **Hyperparameter Tuning of BioMapAI**. We conducted a systematic hyperparameter tuning procedure to optimize BioMapAI’s performance on twelve symptom-specific clinical outcomes and disease status (ME/CFS vs. control). Our goal was to balance predictive accuracy, model complexity, and generalizability across high-dimensional ‘omics datasets. The results of our tuning experiments are illustrated in Supplemental Figure 7. We began with a base BioMapAI architecture consisting of two shared hidden layers (each with 128 nodes), no dropout, no L2 penalty, and training for 1000 epochs. We first investigated how varying the number of shared hidden layers (1, 2, 3, or 4) affected both clinical score prediction (mean squared error, MSE) and disease classification (accuracy). As shown in Supplemental Figure 7A, two shared hidden layers achieved the best predictive performance. Next, we performed a grid search over learning rates {0.01,0.001,0.0005,0.0001,0.00005,0.00001} and batch sizes {32,64,128}. We trained each configuration for 1000 epochs using the Adam optimizer. Supplemental Figure 7B (heatmaps) displays the MSE for each of the 12 clinical scores at different combinations of learning rate and batch size. A learning rate of 0.0005 and batch size 64 emerged as the optimal balance, yielding stable training curves and minimal variance across folds. Although we initially trained for 1000 epochs, we observed that validation metrics consistently stabilized by around 500 epochs. To prevent overfitting and reduce computational burden, we introduced early stopping at 500 epochs in subsequent experiments. We then tuned the number of neurons in each of the two shared hidden layers. Configurations tested included {256,128,64,32,16,8} for the first and the second layer. As shown in Supplemental Figure 7C, while the 128–64 setting performed similarly to other higher-width combinations, we observed that 64–32 minimized overfitting risk yet retained robust predictive accuracy. Thus, we selected 64 neurons in the first shared layer and 32 in the second. To further mitigate overfitting in the hidden layers, we examined dropout rates {0.1,0.2,0.5,0.8}. Supplemental Figure 7D demonstrates that 0.5 offered the best overall balance. We therefore used a 50% dropout after each shared layer. Lastly, we tested L2 penalty strengths λ ∈ {0.1,0.05,0.01,0.005,0.008,0.001,0.0001}. A moderate penalty of λ = 0.008 was selected (Supplemental Figure 7E). Our final chosen hyperparameters include: Two shared hidden layers with sizes 64 and 32, each followed by a ReLU activation and 50% dropout; Batch size = 64, 500 epochs with early stopping; An Adam optimizer (initial learning rate = 0.0005, β1 = 0.9, β2 = 0.999, ε = 1 × 10 − 7), L2 penalty λ = 0.008. We observed that the model’s overall performance (MSE on symptom scores, accuracy for ME/CFS classification) was not highly sensitive to small deviations in these hyperparameters. Even with the baseline configuration (128 nodes, no dropout, no penalty), the predictive performance was reasonable; however, this final tuned setup led to an improvement of approximately 5–10% and yielded more stable and generalizable outcomes across the five ‘omics datasets.
5. **Sensitivity Analyses of BioMapAI.** For sensitivity analysis of BioMapAI, we first re-trained our final BioMapAI configuration ten times with different random initializations. Classification metrics and regression metrics (MSE) for the twelve clinical outcomes were collected. As shown in Supplemental Table 3, the standard deviations (SD) were minimal (<5%) across these ten runs, indicating that BioMapAI is robust to changes in random seed initialization. We also evaluated three similarly performing model architectures (chosen based on grid search results) that yield near-identical or slightly different loss values: **Model 1:** 128 nodes in the first shared layer, 32 nodes in the second shared layer, λ = 0.008; **Model 2:** 32 nodes in the first shared layer, 32 nodes in the second shared layer, λ = 0.008; **Model 3:** 64 nodes in the first shared layer, 32 nodes in the second shared layer, λ = 0.005. As shown in Supplemental Table 3, while minor fluctuations in classification performance were observed, the results were genernally consistent. This underscores BioMapAI’s stability: adjusting the number of neurons in the shared layers or slightly altering the L2 penalty does not substantially degrade classification or regression outcomes. Collectively, these analyses confirm that BioMapAI’s core design is not overly sensitive to small architectural or regularization variations. Even when trained with alternative hyperparameter settings, the model yields robust and consistent performance on both classification (ME/CFS vs. control) and symptom severity score learning.
6. **External Validation with Independent Dataset.** To validate BioMapAI’s robustness in binary classification, we utilized 4 external cohorts^25,26,27,28^ comprising more than 100 samples. For these external cohorts, only binary classification is available. A detailed summary of data collection for these cohorts is provided in Supplemental Table 4. For each external cohort, we processed the raw data (if available) using our in-house pipeline. The features in the external datasets were aligned to match those used in BioMapAI by reindexing the datasets. The overlap between the features in the external dataset and BioMapAI’s feature set was calculated to determine feature coverage. Any missing features were imputed with zeros to maintain consistency across datasets. The input data was then standardized as BioMapAI. We loaded the pre-trained BioMapAI, GBDT, and DNN for comparison. LR and SVM were excluded because they did not perform well during the in-cohort training process. The performance of the models was evaluated using the same binary classification evaluation metrics. Evaluation metrics detailed in Supplemental Table 4.

#### 3. BioMapAI Decode Module: SHAP

BioMapAI is designed to be explainable, ensuring that it not only reconstructs and predicts accurately but also is interpretable, which is particularly crucial in the biological domain. To achieve this, we incorporated SHapley Additive exPlanations (SHAP) into our framework. SHAP offers a consistent measure of feature importance by quantifying the contribution of each input feature to the model’s output.^104^

We applied SHAP to BioMapAI to interpret the results, following these three steps:

1. **Model Reconstruction.** BioMapAI’s architecture includes two shared hidden layers - *Z*^1^, *Z*^2^- and one parallel sub-layers - *Z*^3^-for each object *y*_*i*_. To decode the feature contributions for each object *y*_*i*_, we reconstructed sub-models from single comprehensive model:

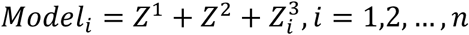

where *n* is the number of learned objects.
2. **SHAP Kernel Explainer.** For each reconstructed model, *Model*_*i*_, we used the SHAP Kernel Explainer to compute the feature contributions. The explainer was initialized with the model’s prediction function and the input data *X*:

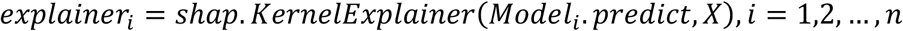

Then SHAP values were computed to determine the contribution of each feature to *y*_*i*_:

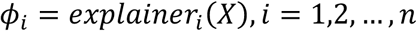

The kernel explainer is a model-agnostic approach that approximates SHAP by evaluating the model with and without the feature of interest and then assigning weights to these evaluations to ensure fairness. For each *model*_*i*_, with each feature *j*:

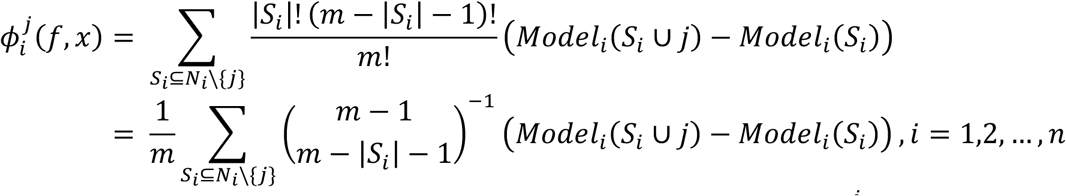

where *n* is the number of learned objects, *m* is the total number of features, Φ_i_^j^ is the Shapley value for feature *j* in *model*_*i*_, *N*_*i*_ is the full set of features in *model*_*i*_, *S*_*i*_ is the subset of features not including feature *j*, *Model*_*i*_(*S*_*i*_) is the model prediction for the subset *S*_*i*_. The SHAP value matrix, Φ_*i*_, were further reshaped to align with the input data dimensions.
3. **Feature Categorization.** Analyzing the SHAP value matrices, [Φ_1_, Φ_2_, …, Φ_*n*_], features can be roughly assigned to two categories: shared features - important to all outputs; or specific features - specifically important to individual outputs. We set the cutoff at 75%, where features consistently identified as top contributors in 75% of the models were classified as shared important features, termed disease-specific biomarkers. Features that were top contributors in only a few models were classified as specific important features, termed symptom-specific biomarkers. By reconstructing individual models, *Model*_*i*_, for each object, *y*_*i*_, and applying SHAP explainer individually, we effectively decoded the contributions of input features to BioMapAI’s predictions. This method allowed us to categorize features into shared and specific categories— termed as disease-specific and symptom-specific biomarkers—providing novel interpretations of the ‘omics feature contribution to clinical symptoms.
4. **Interaction Types of Important Feature. Linear (Monotonic) Relationship**: A feature *x* and a symptom *y* follow a roughly linear (or strictly monotonic) trend when the change in *y* can be approximated by a single slope over *x*’s range. Formally, *y* ≈ α + β*x*, with β ≠ 0, implying a consistently increasing (β > 0) or decreasing (β < 0) trend. Biologically, as the biomarker goes up, the symptom steadily increases (positive β) or decreases (negative β). **Biphasic Relationship.** A biomarker *x* relates to a symptom *y* through a two-phase pattern, such as a U-shaped or inverted U-shaped curve. One way to represent this is by including a squared term: *y* ≈ α + β1*x* + β2*x*^2^, with β2 ≠ 0. Biologically, this often reveals that both very low and very high biomarker values are associated with greater symptom severity, whereas moderate values relate to reduced severity (or vice versa). **Dispersed Relationship**. If there is no single coherent shape (linear or otherwise) that describes the biomarker–symptom relationship across all individuals. Instead, contributions may appear sparse (affecting only a small subset of participants) or highly variable with no dominant pattern. Biologically, this is a typical relationship at KEGG profile in our case, where different individuals can exhibit different directions or magnitudes of effect, leading to scattered or “patchy” patterns.
5. **Stability of SHAP Values**. To the stability of SHAP values under repeated experiments and similar model configurations, we conducted re-ran the Same Data with the Same Architecture (Different Random Seeds) as above. We then computed the standard deviation (SD) of the SHAP values for each feature. Over 90% of features exhibit less than 3% variation in their SHAP contributions across runs, indicating that the top features remain highly consistent despite random seed variation. We also computed SHAP values for each of the three alternative model architectures (Model 1, Model 2, Model 3) described above. Despite their slight architectural or regularization differences, the top 50 features identified by SHAP largely overlapped with those from the final BioMapAI. While some lower-ranked features did differ across models, those changes accounted for less than 5% of the total SHAP variance, suggesting that the core set of important predictors remains stable. Consequently, the minor variations observed are unlikely to affect clinical interpretation or downstream analyses. In summary, both random initializations and small architectural changes do not substantially alter the SHAP-based feature importance patterns in BioMapAI. The top features remain consistent, reinforcing the reliability and interpretability of our multi-output deep learning framework.

#### 4. Packages and Tools

BioMapAI was constructed by Tensorflow(v2.12.0)105 and Keras(v2.12.0). ML models were from scikit-learn(v 1.1.2)106, Glmnet models were using R package glmnet107(v4.1-4) and caret108(v6.0.93).

#### 5. Usage of BioMapAI

We have included our GitHub README.md file and introduced a Jupyter notebook for user instruction. Because there are limited large-scale multi-‘omics datasets with sufficient matched clinical data for us to test BioMapAI’s generalizability, we have not trained BioMapAI on other disease states. However, BioMapAI’s specialized deep neural network structured with two shared general layers and one outcome-focused parallel layer should be generalizable and scalable to other cohort studies that aim to utilize ‘omics data for a range of outputs (e.g., not just limited to clinical symptoms). For instance, researchers could employ our model to link whole genome sequencing data with blood or protein measurements. Constructed to automatically adapt to any input matrix and any output matrix, BioMapAI defaults to parallelly align specific layers for each output.

### WGCNA and Network Analysis

To identify co-expressed patterns of each ‘omics, we employed the Weighted Gene Co-expression Network Analysis (WGCNA) using the WGCNA109 package in R. The analysis was performed on preprocessed omics data (Methods): species abundances (Feature N=373, Sample N=479) and KEGG gene abundances (Feature N=4462, Sample N=479) from the microbiome, plasma metabolome (Feature N=395, Sample N=414), immune profiling (Feature N=311, Sample N=489). Network construction and module detection involved choosing soft-thresholding powers tailored to each dataset: 6 for species, 7 for KEGG, 5 for immune, and 6 for metabolomic. The adjacency matrices were transformed into topological overlap matrices (TOM) to reduce noise and spurious associations. Hierarchical clustering was performed using the TOM, and modules were identified using the dynamic tree cut method with a minimum module size of 30 genes. Module eigengenes were calculated, and modules with highly similar eigengenes (correlation > 0.75) were merged. Module-trait relationships were assessed by correlating module eigengenes with clinical traits, and gene significance (GS) and module membership (MM) were used to identify hub genes within significant modules.

Network analysis was conducted using igraph110 in R. Module eigengenes from the WGCNA analysis were extracted for each dataset. A combined network was constructed by calculating Spearman correlation coefficients (corrected, Methods) between the module eigengenes of different datasets, and an adjacency matrix was created based on a threshold of 0.3 (absolute value) to include only significant associations. Network nodes represented module eigengenes and edges represented significant correlations. Degree centrality and betweenness centrality were calculated to identify highly connected and influential nodes. Networks in patient subgroups were displayed as the correlation differences from their healthy counterparts to exclude the influence of covariates. For example, correlations in female patients were compared with female healthy, and correlations in older patients were compared with older healthy.

### Statistical Analysis

The dimensionality reduction analysis was conducted by Principal Correspondence Analysis (PCoA) using sklearn.manifold.MDS function for ‘omics. For combined ‘omics data, PCoA was applied to combined module eigengenes from WGCNA. Fold change of species, genes, immune cells, and metabolites were compared between patient and control groups, short-term and control groups, and long-term and control groups. P values were computed by Wilcoxon signed-rank test with False Discovery Rate (FDR) correction, adjusted for multiple group comparisons. Spearman’s rank correlation was used to assess correlation covariant. P-values were adjusted using Holm’s method, accounting for multiple group comparisons. P value annotations: ns: p > 0.05, *: 0.01 < p <= 0.05, **: 0.001 < p <= 0.01, ***: p <= 0.001.

### Longitudinal Analysis

To capture statistically meaningful temporal signals, we employed various statistical and modeling methods, accounting for both linear and non-linear trends and intra-individual correlations:

1. **Interquartile Range (IQR) and Intraclass Correlation Coefficient (ICC)**. We initially assessed statistics at different time points by computing the IQR and ICC. Data were standardized to a mean of zero and a standard deviation of one to ensure comparability across features with different scales. The IQR quantified variability, while the ICC assessed the dependence of repeated measurements111, indicating the similarity of measurements over time. Data showed no statistical dependence and no trend of stable variance across time points.
2. **Generalized Linear Models (GLMs)**. GLMs112 were then used to analyze the effects of time points, considering age, gender, and their interactions. Time points were included as predictors to reveal changes in dependent variables over time, with interaction terms exploring variations based on age and gender. Random effects accounted for intra-individual correlations. Although 12 features out of 5000 showed weak trends over time (slopes < 0.2), they were not deemed sufficient to be potential longitudinal biomarkers, possibly due to individualized patterns.
3. **Repeated Measures Correlation (rmcorr)**. To better consider individual effects, we employed rmcorr113 to assess consistent patterns of association within individuals over time. This method captured stable within-individual associations across different time points. However, only 30 features out of 5000 showed weak slopes (< 0.3), and these were not considered sufficient to conclude the presence of longitudinal signals.
4. **Smoothing Spline ANOVA (SS-ANOVA)**. We then considered the longitudinal trends could be non-linear and more complex. To model complex, non-linear relationships between response variables and predictors over time, SS-ANOVA114 was used. SS-ANOVA uncovered non-linear trends and interactions in the omics data, however, no strong temporal signals were identified. In conclusion, robust analysis of the longitudinal data, accounting for both linear and non-linear trends and intra-individual correlations, revealed the difficulty in extracting strong and statistically meaningful temporal signals. As Myalgic Encephalomyelitis/Chronic Fatigue Syndrome (ME/CFS) is a disease that usually lasts for decades with non-linear progression, the four-year tracking period with annual measurements is likely insufficient for capturing consistent temporal signals, necessitating longer follow-up periods.

## Data and Code

Metagenomics data is being deposited under the BioProject submission number SUB14546737 and will be publicly available as of the date of publication. Accession numbers are listed in the key resources table. BioMapAI framework is available at https://github.com/ohlab/BioMapAI/codes/AI. All original code, analyzed data and trained model has been deposited at https://github.com/ohlab/BioMapAI. All other ‘omics data, including clinical metadata, are available in Supplementary Tables, GitHub and at the MapMECFS portal (https://mecfs.rti.org/research/). Any additional information required to reanalyze the data reported in this paper is available from the lead contact upon request.

## Acknowledgements and Funding

We are thankful to the Oh, Unutmaz, and Li laboratories for inspiring discussions and acknowledge the contribution of the Genome Technologies Service at The Jackson Laboratory for expert assistance with sample sequencing for the work described in this publication. We also thank the clinical support team at the Bateman Horne Center and all the individuals who participated in this study. This work was funded by 1U54NS105539. JO is additionally supported by the NIH (1 R01 AR078634-01, DP2 GM126893-01, 1 U19 AI142733, 1 R21 AR075174).

## Author Contributions

Conceptualization: DU, JO, SDV, LB, RX; Data Curation: RX, CG, SDV, LB; Formal Analysis: RX; Funding Acquisition: DU, JO, SDV, LB; Clinical sample design and collection: SDV, LB; Investigation: RX, CG, EF, SDV, LB; Project Administration: JO, DU, LB, SDV, CG; Resources: DU, JO, SDV, LB; Supervision: JO; Visualization and Writing: RX, JO; Writing - Review and Editing: RX, CG, SDV, LB, DU, JO.

## Competing Interests

Dr. Suzanne D. Vernon is affiliated and has a financial interest with The BioCollective, a company that provided the BioCollector, the collection kit used for at home stool collection discussed in this manuscript. No other authors have competing interests.

## Lead Contact

Further information and requests for resources and reagents should be directed to the lead contact, Julia Oh.

## Supplemental Figure

**Supplemental Figure 1:**
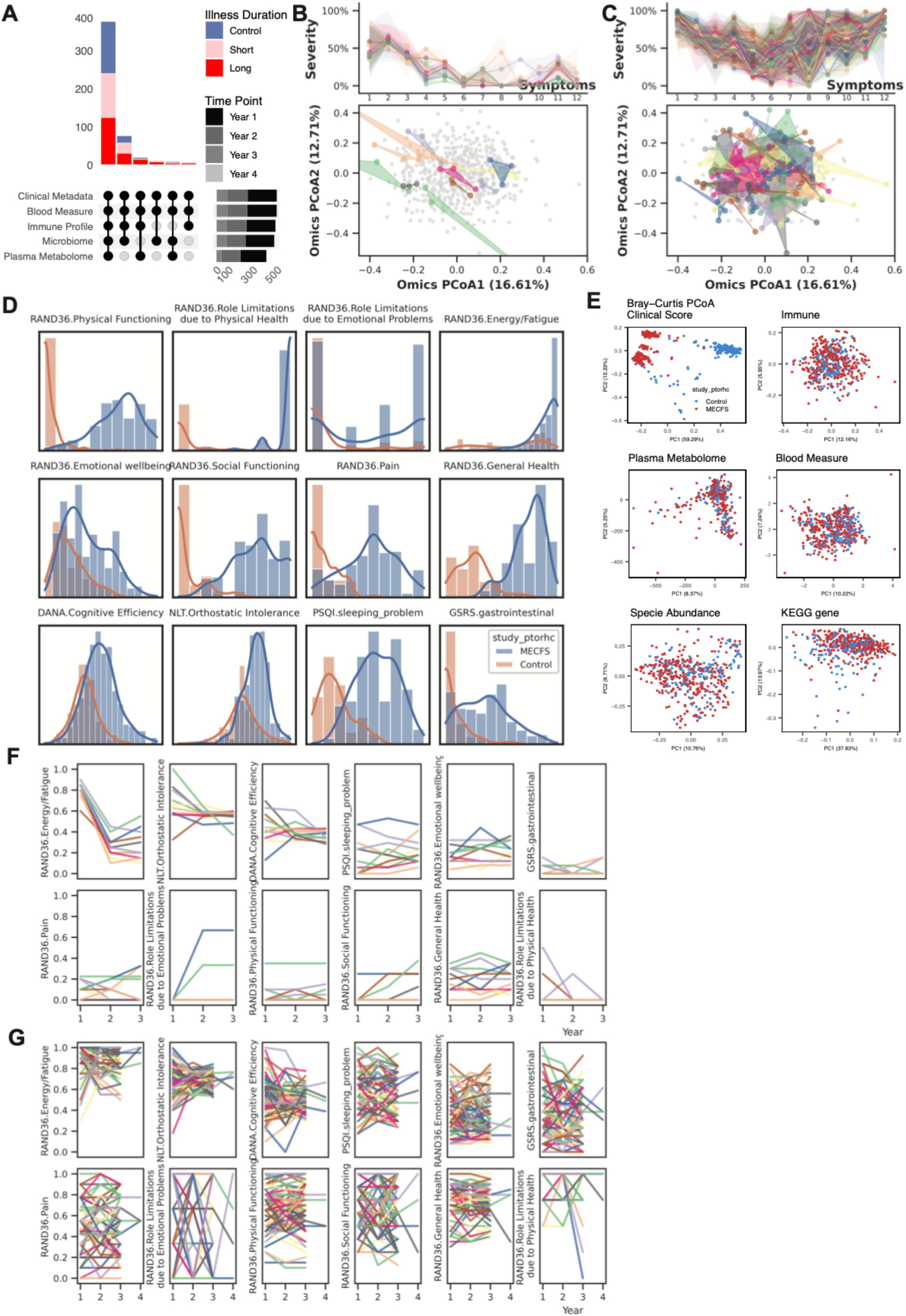
Data Pairedness Overview and Heterogeneity in Healthy and Patients. A) Cohort Composition and Data Collection. Over four years, 515 time points were collected: baseline year from all 249 donors (Healthy N=96, ME/CFS N=153); second year from 168 individuals (Healthy N=58, ME/CFS N=110); third year from 94 individuals (Healthy N=13, ME/CFS N=81); fourth year from N=4 ME/CFS patients. Nearly 400 collection points included complete sets of 5 ‘omics datasets, with others capturing 3-4 ‘omics profiles. Clinical metadata and blood measures were collected at all 515 points. Immune profiles from PBMCs were recorded at 489 points, microbiome data from stool samples at 479 points, and plasma metabolome data at 414 points. A total of 1,471 biosamples were collected. **B-C) Heterogeneity of B) Healthy Controls and C) All Patients in Symptom Severity and ‘Omics Profiles.** Supplemental information for Figure 1B, which shows examples from 20 patients. Variability in symptom severity (top) and ‘omics profiles (bottom) for all healthy controls and all patients with 3-4 time points. The top x-axis numbers represent 12 symptoms, arranged in the same order as Supplemental Figure F-G (left to right, top to bottom). **D) Distribution of 12 Clinical Symptoms in ME/CFS and Control.** Density plots compare the distributions of 12 clinical scores between control (blue) and ME/CFS patients (orange) with the x-axis represents the values of symptom severity (scaled from 0%, no symptom, to 100%, most severe)n and the y-axis represents the frequency (count) of data points. Clinical scores include RAND36 subscales (e.g., Physical Functioning, Emotional Wellbeing), Cognitive Efficiency from the DANA test, Orthostatic Intolerance from the NLT test, Sleep Problems from the PSQI questionnaire, and Gastrointestinal Symptoms from the GSRS questionnaire. **E) Principal Coordinates Analysis (PCoA) of each ‘Omics.** PCoA based on Bray-Curtis distance for clinical scores, immune profiles, plasma metabolome, blood measures, species abundance, and KEGG gene data. Control samples (blue) and ME/CFS patients (red) show distinct clustering. Here, except for the clinical scores, controls are indistinguishable from patients, highlighting the difficulty of building classification models. **F-G) Symptom Progression Over Time in F) Healthy vs. G) ME/CFS Patients**. Symptom progression for each individual (represented by different colors) is shown using line plots of symptom severity (y-axis) over time points (years 1–4). Compared to healthy controls, ME/CFS patients exhibit higher severity (indicated by higher y-axis values), greater heterogeneity (indicated by differences within the patient group), and inconsistent or non-linear progression (indicated by substantial variation over time without a consistent pattern) in clinical symptoms. **Abbreviations:** ME/CFS, Myalgic Encephalomyelitis/Chronic Fatigue Syndrome; PCoA, Principal Coordinates Analysis; RAND36, 36-Item Short Form Health Survey; DANA, DANA Brain Vital; NLT, NASA Lean Test; PSQI, Pittsburgh Sleep Quality Index; GSRS, Gastrointestinal Symptom Rating Scale; KEGG, Kyoto Encyclopedia of Genes and Genomes. **Related to:** Figure 1-2.

**Supplemental Figure 2:**
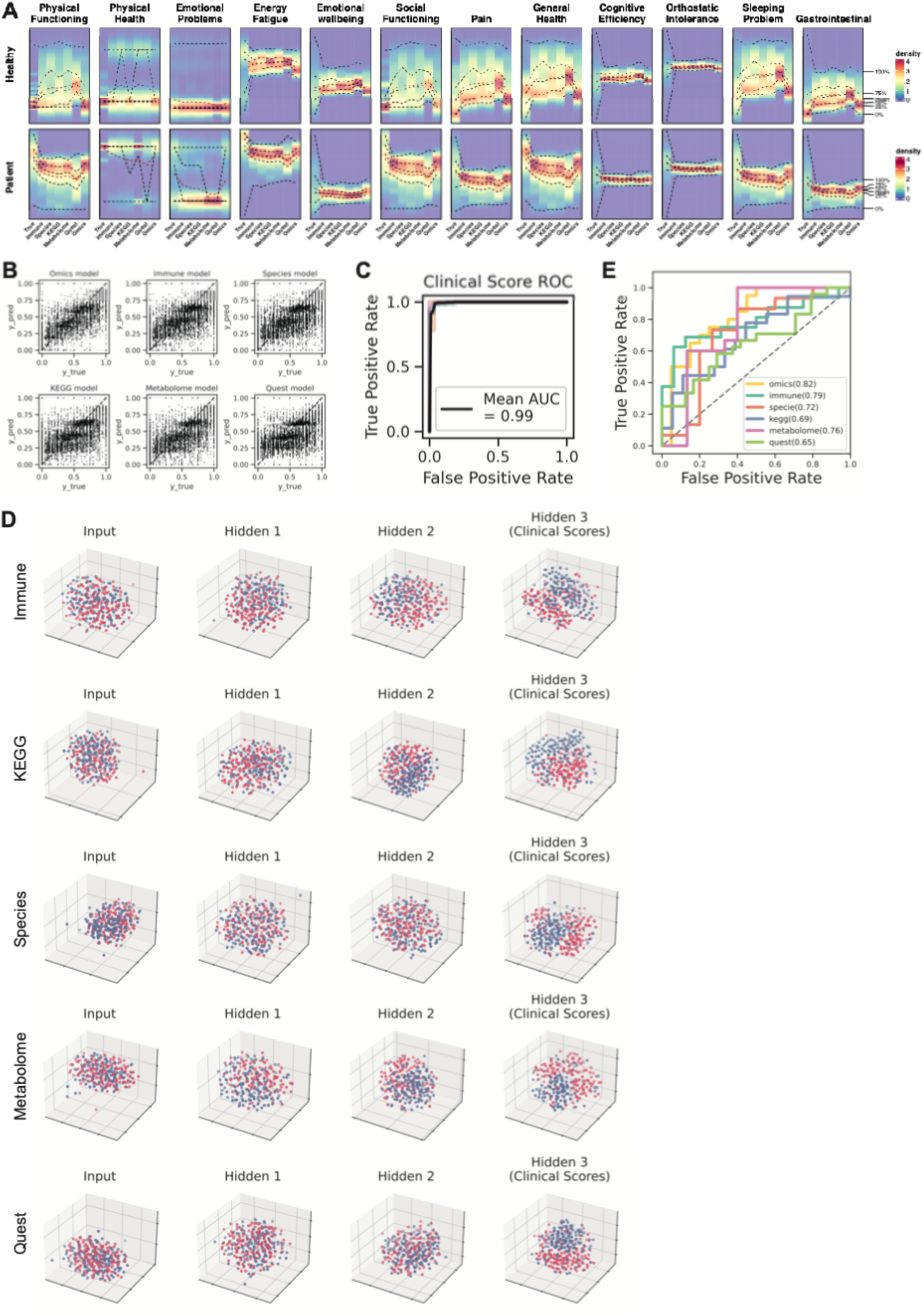
BioMapAI’s Performance at Clinical Score Reconstruction and Disease Classification. A) Density map of True vs. Predicted Clinical Scores. Supplemental information for Figure 2B, which shows three examples. Here, the full set of 12 clinical scores compares the true score, *y* (Column 1), against BioMapAI’s predictions generated from different ‘omics profiles – ŷ_*immune*_, ŷ_*KEGG*_, ŷ_*KEGG*_, ŷ_*metabolome*_, ŷ_*speices*_, ŷ_*quest*_ (Columns 2-7). B) Scatter Plot of True vs. Predicted Clinical Scores. Scatter plots display the relationship between true clinical scores (x-axis) and predicted clinical scores (y-axis) for six different models: Omics, Immune, Species, KEGG, Metabolome, and Quest Labs. Each plot demonstrates the clinical score prediction accuracy for each model. C) ROC Curve for Disease Classification with Original Clinical Scores. The Receiver Operating Characteristic (ROC) curve evaluates the performance of disease classification using the original 12 clinical scores. The mean Area Under the Curve (AUC) is 0.99, indicating high prediction accuracy, which aligns with the clinical diagnosis of ME/CFS based on key symptoms. D) 3D t-SNE Visualization of Hidden Layers. 3D t-SNE plots show how BioMapAI progressively distinguishes disease from control across hidden layers for five trained ‘omics models: Immune, KEGG, Species, Metabolome, and Quest Labs. Each plot uses the first three principal components to show the spatial distribution of control samples (blue) and ME/CFS patients (red). The progression from the input layer (mixed groups) to Hidden Layer 3 (fully separated groups) illustrates how BioMapAI progressively learns to separate ME/CFS from healthy controls. E) ROC Curve for Disease Classification with Held-out Data. ROC curves show BioMapAI’s performance in disease classification with held-out data. Abbreviations: ROC, Receiver Operating Characteristic; AUC, Area Under the Curve; t-SNE, t-Distributed Stochastic Neighbor Embedding; PCs, Principal Components; *y*, True Score; *y*2, Predicted Score. Related to: Figure 2.

**Supplemental Figure 3:**
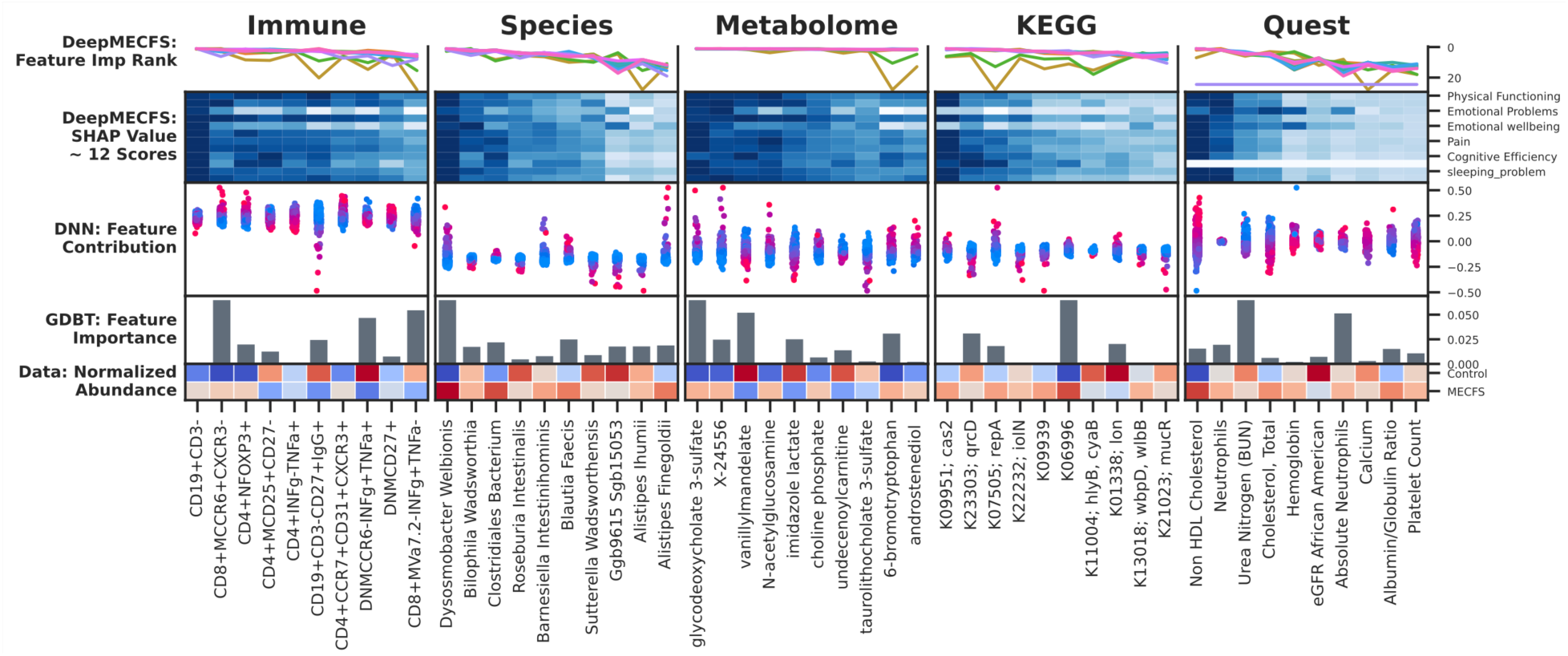
Disease-Specific Biomarkers - Top 10 Biomarkers Shared across Clinical Symptoms and Multiple Models. Through the top 30 high-ranking features for each score, we discovered that the most critical features for all 12 symptoms were largely shared and consistently validated across ML and DL models, particularly the foremost 10. Here, this multi-panel figure presents the top 10 most significant features identified by BioMapAI across five ‘omics profiles, highlighting their importance in predicting clinical symptoms and diagnostic outcomes across BioMapAI, DNN, and GBDT models, along with their data prevalence. Each vertical section represents one ‘omics profile, with columns of biomarkers ordered by average feature importance from right to left. From top to bottom: *1. Feature Importance Ranking in BioMapAI.* Lines depict the rank of feature importance for each clinical score, color-coded by the 12 clinical scores. Consistency among the top 5 features suggests they are shared disease biomarkers crucial for all clinical symptoms; *2. Heatmap of SHAP Values from BioMapAI.* This heatmap shows averaged SHAP values with the 12 scores on the rows and the top 10 features in the columns. Darker colors indicate a stronger impact on the model’s output; *3. Swarm Plot of SHAP Values from DNN.* This plot represents the distribution of feature contributions from DNN, which is structurally similar to BioMapAI but omits the third hidden layer (*Z*^3^). SHAP values are plotted vertically, ranging from negative to positive, showing each feature’s influence on prediction outcomes. Points represent individual samples, with color gradients denoting actual feature values. For instance, *Dysosmobacteria welbionis*, identified as the most critical species, shows that greater species relative abundance correlates with a higher likelihood of disease prediction; *4. Bar Graphs of Feature Importance in GBDT*. GBDT is another machine learning model used for comparison. Each bar’s height indicates a feature’s significance within the GBDT model, providing another perspective on the predictive relevance of each biomarker; *5. Heatmap of Normalized Raw Abundance Data.* This heatmap compares biomarker prevalence between healthy and disease states, with colors representing z-scored abundance values, highlighting biomarker differences between groups. **Abbreviations:** DNN: Here refer to our deep Learning model without the hidden 3, ‘spread out’ layer; GBDT: Gradient Boosting Decision Tree; SHAP: SHapley Additive exPlanations. **Supporting Materials:** Supplemental Table 5. **Related to:** Figure 3.

**Supplemental Figure 4:**
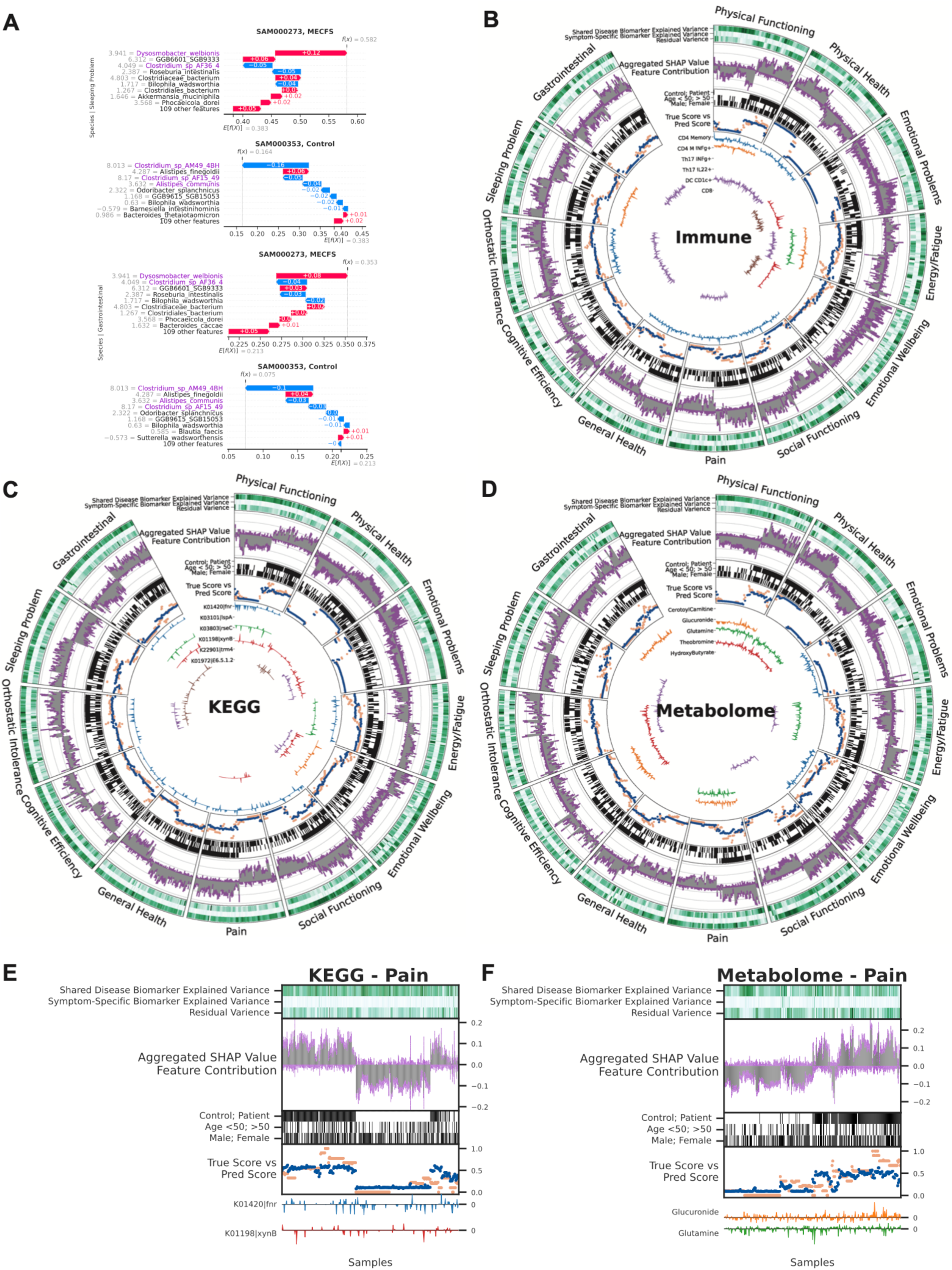
Symptom-Specific Biomarkers - Immune, KEGG and Metabolome Models. By linking ‘omics profiles to clinical symptoms, BioMapAI identified unique symptom-specific biomarkers in addition to disease-specific biomarkers (Supplemental Figure 3). Each ‘omics has a circularized diagram (Figure 3A, Supplemental Figure 4B-D) to display how BioMapAI use this ‘omics profile to predict 12 clinical symptoms and to discuss the contribution of disease- and symptom-specific biomarkers. Detailed correlation between symptom-specific biomarkers and their corresponding symptoms is in Supplemental Figure 5. **A) Examples of Sleeping Problem-Specific Species’ and Gastrointestinal-Specific Species’ Contributions.** Supplemental information for Figure 3D, which shows the contribution of pain-specific species. **B-D) Circularized Diagram for Immune, KEGG and Metabolome Models.** Supplemental information for Figure 3A, which shows the species model. **E-F) Zoomed Segment for Pain in KEGG and Metabolome Model.** Supplemental information for Figure 3B, which shows the zoomed segment for pain in the species and immune models. *Note, the reported biomarkers were calculated using the entire dataset and were not validated on held-out data. **Abbreviations and Supporting Materials:** Supplemental Figure 5. **Related to:** Figure 3.

**Supplemental Figure 5:**
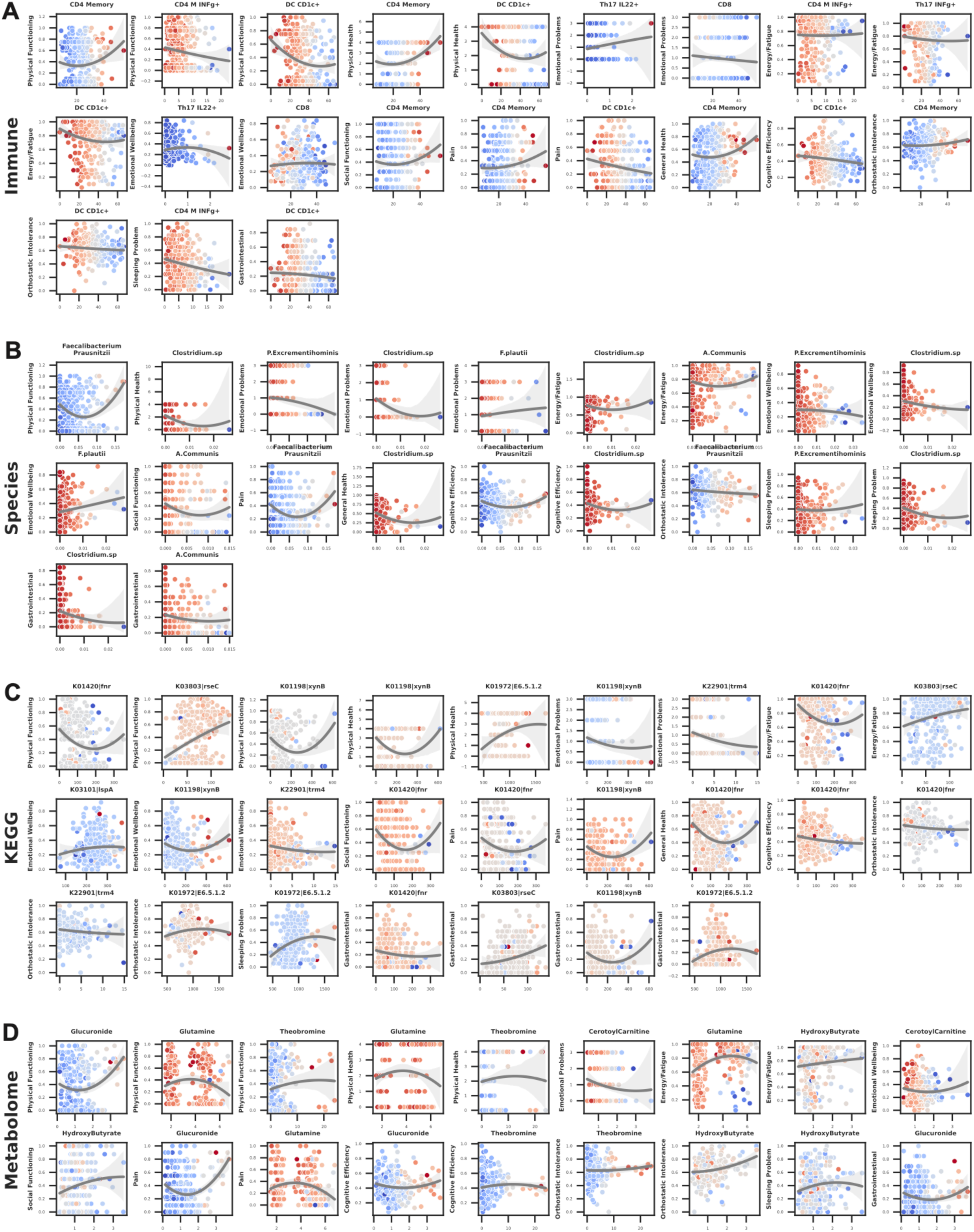
Symptom-Specific Biomarkers - Different Correlation Patterns of Biomarkers to Symptom. Supplemental information for Figure 3C, which shows six pain biomarkers from multiple models. Here for each ‘omics, we plotted the correlation of symptom-specific biomarkers (x-axis) to its related symptom (y-axis), colored by SHAP value (contribution to the symptom). **Abbreviations:** CD4, Cluster of Differentiation 4; CD8, Cluster of Differentiation 8; IFNg, Interferon Gamma; DC, Dendritic Cells; MAIT, Mucosal-Associated Invariant T; Th17, T helper 17 cells; CD4+ TCM, CD4+ Central Memory T cells; DC CD1c+ mBtp+, Dendritic Cells expressing CD1c+ and myelin basic protein; DC CD1c+ mHsp, Dendritic Cells expressing CD1c+ and heat shock protein; CD4+ TEM, CD4+ Effector Memory T cells; CD4+ Th17 rfx4+, CD4+ T helper 17 cells expressing RFX4; *F. prausnitzii, Faecalibacterium prausnitzii; A. communis, Akkermansia communis*; NAD, Nicotinamide Adenine Dinucleotide. **Related to:** Figure 3.

**Supplemental Figure 6:**
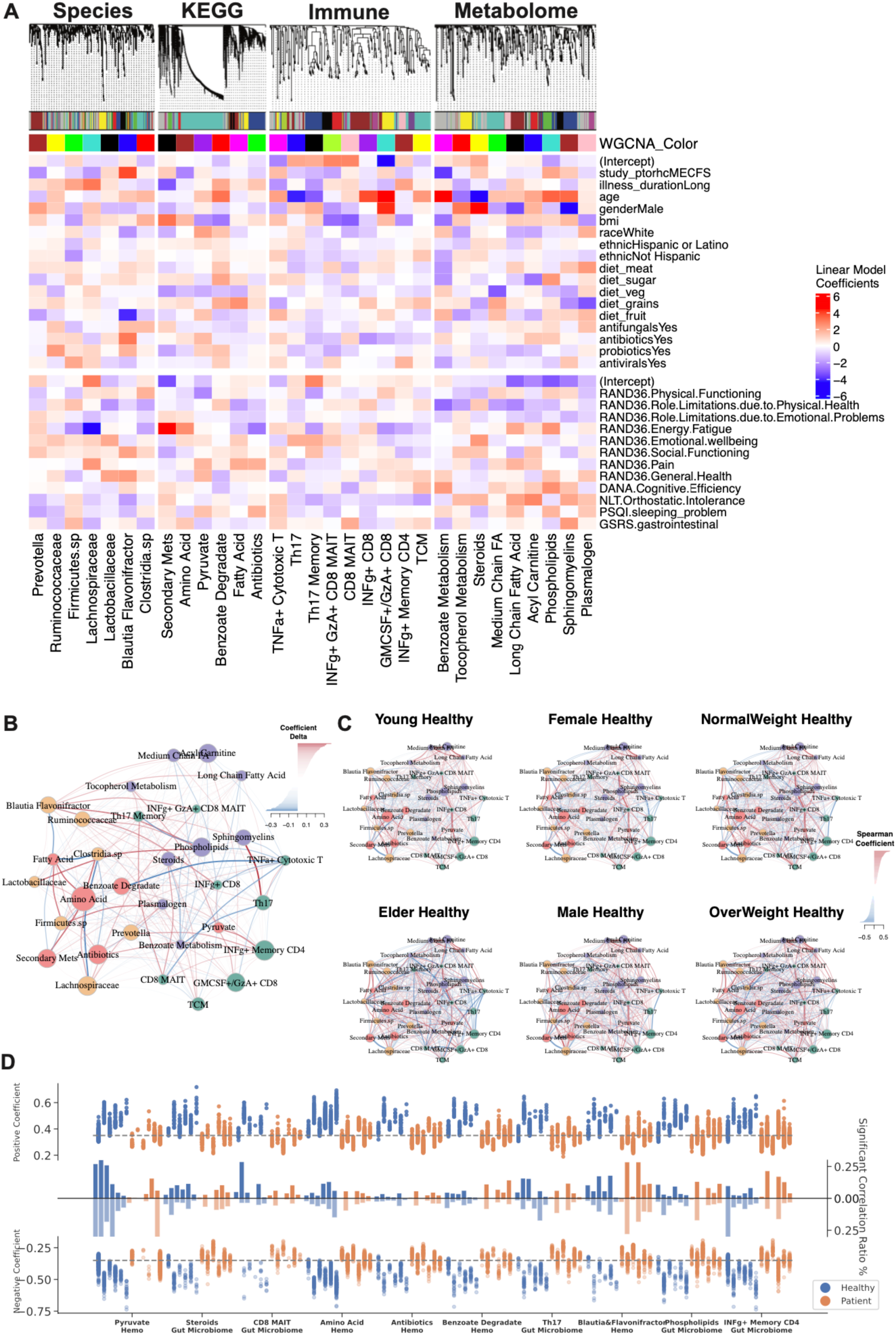
‘Omics WGCNA Modules and Host-Microbiome Network. A) Correlation of WGCNA Modules with Clinical Metadata. Weighted Gene Co-expression Network Analysis (WGCNA) was used to identify co-expression modules for each ‘omics layer: species, KEGG, immune, and metabolome. The top dendrograms show hierarchical clustering of ‘omics features, with modules identified. The bottom heatmap shows the relationship of module eigengenes (colored as per dendrogram) with clinical metadata – including demographic information and environmental factors - and 12 clinical scores. General linear models were used to determine the primary clinical drivers for each module, with the color gradient representing the coefficients (red = positive, blue = negative). Microbial modules were influenced by disease presence and energy-fatigue levels, while metabolome and immune modules correlated with age and gender. **B-C) Microbiome-Immune-Metabolome Network in B) Patient and C) Healthy Subgroups.** Supplemental information for Figure 4A (Healthy Network) and 4B (Patient Subgroups). Figure 4A is the healthy network; here, Supplemental Figure 6B presented the shifted correlations in all patients. Figure 4B represented the network in patient subgroups; here, Supplemental Figure 6C is the corresponding healthy counterpart, for example, female patients were compared with female controls to exclude gender influences. **D) Differences in Host-Microbiome Correlations between Healthy and Patient Subgroups.** Selected host-microbiome module pairs are grouped on the x-axis (e.g., pyruvate to blood modules, steroids to gut microbiome). Significant positive and negative correlations (top and bottom y-axis) of module members pairs are shown as dots for each subgroup (blue = healthy, orange = patient) (Spearman, adjusted p < 0.05), from left to right: Young, Elder, Female, Male, NormalWeight, OverWeight Healthy and Young, Elder, Female, Male, NormalWeight, OverWeight Patient. The middle bars represent the total count of associations. This panel highlights the shifts in host-microbiome networks from health to disease, for example, in patients, the loss of pyruvate to host blood modules correlation and the increase of INFg+ CD4 memory correlation with gut microbiome. **Abbreviations:** WGCNA, Weighted Gene Co-expression Network Analysis; AA, Amino Acids; SCFA, Short-Chain Fatty Acids; IL, Interleukin; GM-CSF, Granulocyte-Macrophage Colony-Stimulating Factor. **Related to**: Figure 4.

**Supplemental Figure 7:**
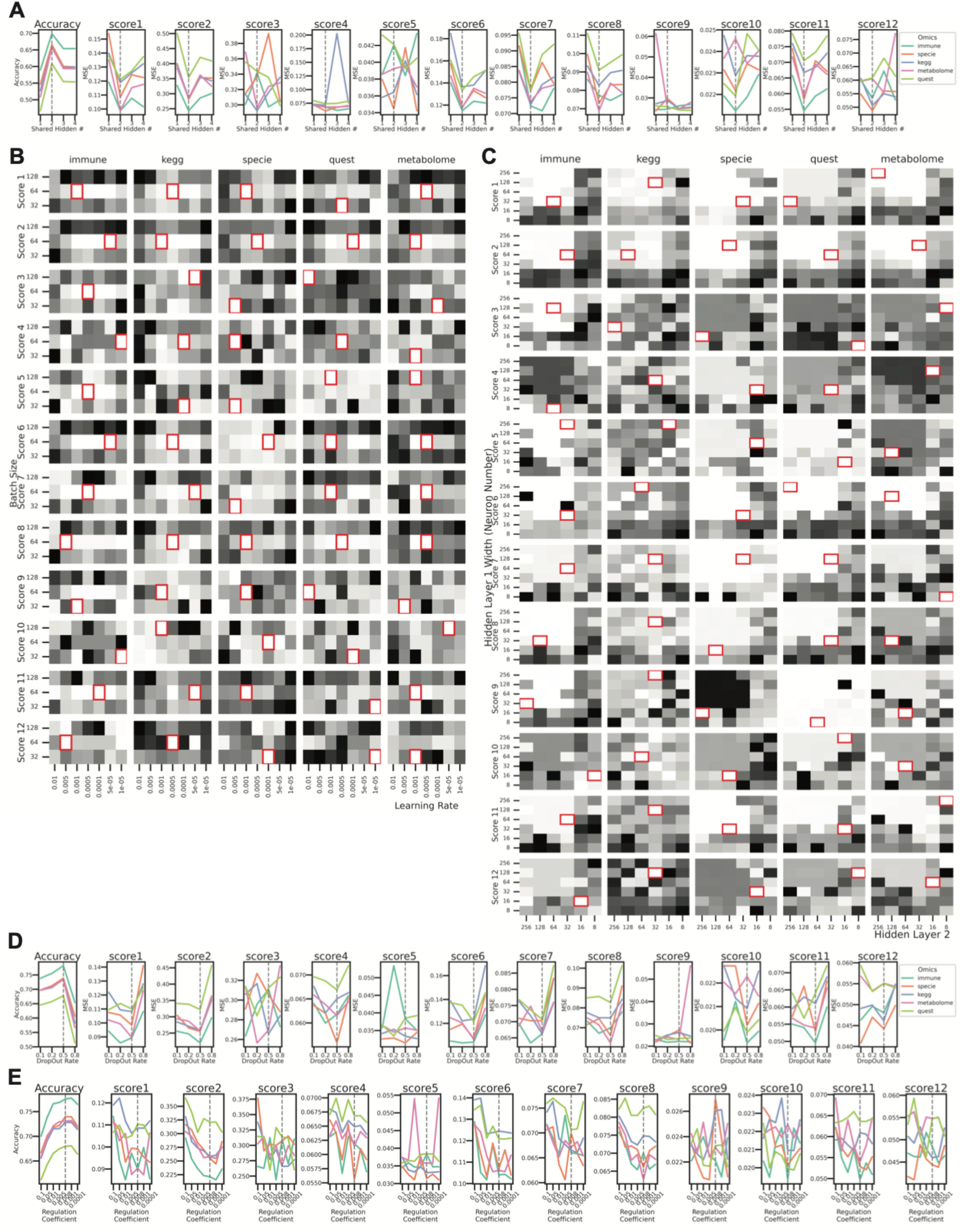
Hyperparameter Tuning of BioMapAI. This figure illustrates how BioMapAI’s predictive performance for 12 symptom-specific clinical objectives and disease classification (ME/CFS vs. control) responds to different hyperparameter settings across five ‘omics datasets (species abundance, KEGG gene abundances, plasma metabolome, immune profiling, and quest blood measurements). Each sub-panel shows a comparison of performance metrics (e.g., mean squared error for clinical scores, classification accuracy for disease classification) versus the tested hyperparameter values. For mean squared error (MSE) of clinical scores, smaller values indicate better predictions, as predicted scores are closer to true scores. For classification accuracy, larger values reflect better performance. **A) Number of Shared Hidden Layers.** The y-axis represents performance metrics tuned against the number of shared hidden layers {1,2,3,4} on the x-axis Each ‘omics dataset is distinguished by color. Two shared hidden layers were selected, as this configuration demonstrated the best balance between predictive accuracy for clinical scores and disease classification. **B) Grid Search for Learning Rate and Batch Size.** Each heatmap represents one ‘omics dataset (columns), with rows corresponding to individual clinical scores (total of 12). Colors indicate MSE between predicted *y*_*pred*_ and *y*_*true*_ values across combinations of learning rates {0.01,0.001,0.0005,0.0001,0.00005,0.00001} and batch sizes {32,64,128}. Darker colors represent higher MSE (worse prediction), while lighter colors indicate lower MSE (better prediction). Red boxes mark optimal settings. A learning rate of 0.0005 and batch size of 64 achieved stable training with minimal variance across predictions. **C) Grid Search for Number of Neurons in Each Shared Layer.** Similar to (B), this panel visualizes the tuning of network width for each shared layer. Configurations tested include {256,128,64,32,16,8}. While configurations like 128–64 (Layer 1) and 64–32 (Layer 2) performed similarly, 64–32 was chosen for minimizing overfitting while preserving predictive accuracy. **D) Dropout Rate.** The x-axis shows tested dropout rates {0.1,0.2,0.5,0.8}, while the y-axis tracks performance metrics as in (A). A dropout rate of 50% (0.5) provided the best trade-off between overfitting control and prediction stability. **D) L2 Penalty Rate.** Each line or bar corresponds to different regularization strengths λ ∈ {0.1,0.05,0.01,0.005,0.008,0.001,0.0001}. A moderate penalty of λ = 0.008 was selected, offering an optimal balance between overfitting prevention and model capacity. Together, these panels demonstrate how each hyperparameter affects BioMapAI’s ability to predict 12 clinical scores and classify disease status across five distinct ‘omics datasets. The final configuration— two shared hidden layers (64 and 32 neurons), a learning rate of 0.0005, batch size of 64, dropout of 50%, and L2 penalty λ = 0.008 —achieved optimal balance between predictive performance and generalizability for high-dimensional ‘omics data. **Abbreviations: ME/CFS:** Myalgic Encephalomyelitis/Chronic Fatigue Syndrome; **KEGG:** Kyoto Encyclopedia of Genes and Genomes**; MSE:** Mean Squared Error. **Related to:** Methods.

## Supplemental Table

**Supplemental Table 1** Sample Metadata and Clinical Scores

**Supplemental Table 2** Model Performance at Reconstructing Twelve Clinical Scores: Averaged Average Mean Squared Error by Model and Model Sensitivity Analysis

**Supplemental Table 3** Model Performance in Diagnostic Comparison—Within-Cohort, Cross-Validated and Held-Out by Various ML and DL Models

**Supplemental Table 4** Model Performance in Diagnostic Comparison—Across Independent Cohorts

**Supplemental Table 5** Disease-Specific Biomarker: Averaged Feature Contribution of BioMapAI, DNN and GDBT

**Supplemental Table 6** Symptom-Specific Biomarker: Distinct Sets of Biomarkers for Each Symptom

**Supplemental Table 7** WGCNA Module Eigengene

**Supplemental Table 8** Targeted Pathways: Normalized Gene Read Counts and Their Correlation with Blood Responders

## Notes

### Summary of Updates

Reclaimed the value of the study and improved the algorithm

## References

1. Cortes Rivera, M., Mastronardi, C., Silva-Aldana, C. T., Arcos-Burgos, M. C, Lidbury, B. A. Myalgic Encephalomyelitis/Chronic Fatigue Syndrome: A Comprehensive Review. Diagnostics G, 91 (2019).

2. Sweetman, E. et al. Current Research Provides Insight into the Biological Basis and Diagnostic Potential for Myalgic Encephalomyelitis/Chronic Fatigue Syndrome (ME/CFS). Diagnostics G, 73 (2019).

3. Noor, N. et al. A Comprehensive Update of the Current Understanding of Chronic Fatigue Syndrome. *Anesthesiol*. Pain Med. 11, e113629 (2021).

4. Ruiz-Pablos, M., Paiva, B., Montero-Mateo, R., Garcia, N. C, Zabaleta, A. Epstein-Barr Virus and the Origin of Myalgic Encephalomyelitis or Chronic Fatigue Syndrome. Front. Immunol. 12, 656797 (2021).

5. Su, R. et al. The TLR3/IRF1/Type III IFN Axis Facilitates Antiviral Responses against Enterovirus Infections in the Intestine. mBio 11, 10.1128/mbio.02540-20 (2020).

6. Anderson, D. E. et al. Lack of cross-neutralization by SARS patient sera towards SARS-CoV-2. Emerg. Microbes Infect. 9, 900–902 (2020).

7. Cairns, R. C, Hotopf, M. A systematic review describing the prognosis of chronic fatigue syndrome. Occup. Med. Oxf. Engl. 55, 20–31 (2005).

8. de Mel, S., Lim, S. H., Tung, M. L. C, Chng, W.-J. Implications of Heterogeneity in Multiple Myeloma. BioMed Res. Int. 2014, 232546 (2014).

9. Wallstrom, G., Anderson, K. S. C, LaBaer, J. Biomarker Discovery for Heterogeneous Diseases.

10. Weyand, C. M., McCarthy, T. G. C, Goronzy, J. J. Correlation between disease phenotype and genetic heterogeneity in rheumatoid arthritis. J. Clin. Invest. 95, 2120–2126 (1995).

11. Poenaru, S., Abdallah, S. J., Corrales-Medina, V. C, Cowan, J. COVID-19 and post-infectious myalgic encephalomyelitis/chronic fatigue syndrome: a narrative review. Ther. Adv. Infect. Dis. 8, 20499361211009385 (2021).

12. Reuken, P. A. et al. Longterm course of neuropsychological symptoms and ME/CFS after SARS-CoV-2-infection: a prospective registry study. Eur. Arch. Psychiatry Clin. Neurosci. (2023) doi:10.1007/s00406-023-01661-3.

13. Hare, P. J., LaGree, T. J., Byrd, B. A., DeMarco, A. M. C, Mok, W. W. K. Single-Cell Technologies to Study Phenotypic Heterogeneity and Bacterial Persisters. Microorganisms 9, 2277 (2021).

14. Cohen, R. M., Haggerty, S. C, Herman, W. H. HbA1c for the Diagnosis of Diabetes and Prediabetes: Is It Time for a Mid-Course Correction? J. Clin. Endocrinol. Metab. 95, 5203– 5206 (2010).

15. Zhou, W. et al. Longitudinal multi-omics of host–microbe dynamics in prediabetes. Nature 569, 663–671 (2019).

16. Hong, S. et al. Cancer Statistics in Korea: Incidence, Mortality, Survival, and Prevalence in 2017. Cancer Res. Treat. 52, 335–350 (2020).

17. Zeeshan, S., Xiong, R., Liang, B. T. C, Ahmed, Z. 100 years of evolving gene–disease complexities and scientific debutants. Brief. Bioinform. 21, 885–905 (2020).

18. Bretherick, A. D. et al. Typing myalgic encephalomyelitis by infection at onset: A DecodeME study. NIHR Open Res. 3, 20 (2023).

19. Bae, J. C, Lin, J.-M. S. Healthcare Utilization in Myalgic Encephalomyelitis/Chronic Fatigue Syndrome (ME/CFS): Analysis of US Ambulatory Healthcare Data, 2000–2009. Front. Pediatr. 7, (2019).

20. Zheng, Y. C, Zhu, Z. Editorial: Retrieving meaningful patterns from big biomedical data with machine learning approaches. Front. Genet. 14, (2023).

21. Leelatian, N. et al. Unsupervised machine learning reveals risk stratifying glioblastoma tumor cells. eLife 9, e56879 (2020).

22. Su, Q., et al. The gut microbiome associates with phenotypic manifestations of post-acute COVID-19 syndrome. Cell Host Microbe 32, 651–660.e4 (2024).

23. Bourgonje, A. R., van Goor, H., Faber, K. N. C, Dijkstra, G. Clinical Value of Multiomics-Based Biomarker Signatures in Inflammatory Bowel Diseases: Challenges and Opportunities. Clin. Transl. Gastroenterol. 14, e00579 (2023).

24. Marcos-Zambrano, L. J. et al. Applications of Machine Learning in Human Microbiome Studies: A Review on Feature Selection, Biomarker Identification, Disease Prediction and Treatment. Front. Microbiol. 12, (2021).

25. Guo, C. et al. Deficient butyrate-producing capacity in the gut microbiome is associated with bacterial network disturbances and fatigue symptoms in ME/CFS. Cell Host Microbe 31, 288–304.e8 (2023).

26. Raijmakers, R. P. H. et al. Multi-omics examination of Q fever fatigue syndrome identifies similarities with chronic fatigue syndrome. J. Transl. Med. 18, 448 (2020).

27. Germain, A., et al. Plasma metabolomics reveals disrupted response and recovery following maximal exercise in myalgic encephalomyelitis/chronic fatigue syndrome. JCI Insight 7, (2023).

28. Che, X., et al. Metabolomic Evidence for Peroxisomal Dysfunction in Myalgic Encephalomyelitis/Chronic Fatigue Syndrome. Int. J. Mol. Sci. 23, 7906 (2022).

29. Liñares-Blanco, J., Fernandez-Lozano, C., Seoane, J. A. C, López-Campos, G. Machine Learning Based Microbiome Signature to Predict Inflammatory Bowel Disease Subtypes. Front. Microbiol. 13, (2022).

30. He, F. et al. Development and External Validation of Machine Learning Models for Diabetic Microvascular Complications: Cross-Sectional Study With Metabolites. J. Med. Internet Res. 26, e41065 (2024).

31. Hawken, S., et al. External validation of machine learning models including newborn metabolomic markers for postnatal gestational age estimation in East and South-East Asian infants. Preprint at 10.12688/gatesopenres.13131.2 (2021).

32. Mora-Ortiz, M., Trichard, M., Oregioni, A. C, Claus, S. P. Thanatometabolomics: introducing NMR-based metabolomics to identify metabolic biomarkers of the time of death. Metabolomics 15, 37 (2019).

33. Balasubramanian, R. et al. Metabolomic profiles associated with all-cause mortality in the Women’s Health Initiative. Int. J. Epidemiol. 49, 289–300 (2020).

34. Li, H., Ren, M. C, Li, Q. 1H NMR-Based Metabolomics Reveals the Intrinsic Interaction of Age, Plasma Signature Metabolites, and Nutrient Intake in the Longevity Population in Guangxi, China. Nutrients 14, 2539 (2022).

35. Kondoh, H. C, Kameda, M. Metabolites in aging and aging-relevant diseases: Frailty, sarcopenia and cognitive decline. Geriatr. Gerontol. Int. 24, 44–48 (2024).

36. Peng, S., Shen, Y., Wang, M. C, Zhang, J. Serum and CSF Metabolites in Stroke-Free Patients Are Associated With Vascular Risk Factors and Cognitive Performance. Front. Aging Neurosci. 12, (2020).

37. Duerler, P., Vollenweider, F. X. C, Preller, K. H. A neurobiological perspective on social influence: Serotonin and social adaptation. J. Neurochem. 162, 60–79 (2022).

38. Pomrenze, M. B., Paliarin, F. C, Maiya, R. Friend of the Devil: Negative Social Influences Driving Substance Use Disorders. Front. Behav. Neurosci. 16, (2022).

39. Laslett, A.-M. Commentary on Bischof, et al.: Empirical and conceptual paradigms for studying secondary impacts of a person’s substance use. Addiction 117, 3148–3149 (2022).

40. Carco, C. et al. Increasing Evidence That Irritable Bowel Syndrome and Functional Gastrointestinal Disorders Have a Microbial Pathogenesis. Front. Cell. Infect. Microbiol. 10, (2020).

41. Saffouri, G. B. et al. Small intestinal microbial dysbiosis underlies symptoms associated with functional gastrointestinal disorders. Nat. Commun. 10, 2012 (2019).

42. Liang, S., Wu, X., Hu, X., Wang, T. C, Jin, F. Recognizing Depression from the Microbiota–Gut–Brain Axis. Int. J. Mol. Sci. 19, 1592 (2018).

43. Zhu, F., Tu, H. C, Chen, T. The Microbiota–Gut–Brain Axis in Depression: The Potential Pathophysiological Mechanisms and Microbiota Combined Antidepression Effect. Nutrients 14, 2081 (2022).

44. Topan, R. C, Scott, S. M. Sleep: An Overlooked Lifestyle Factor in Disorders of Gut-Brain Interaction. Curr. Treat. Options Gastroenterol. 21, 435–446 (2023).

45. Moens de Hase, E., et al. Impact of metformin and Dysosmobacter welbionis on diet-induced obesity and diabetes: from clinical observation to preclinical intervention. Diabetologia 67, 333–345 (2024).

46. Amabebe, E., Robert, F. O., Agbalalah, T. C, Orubu, E. S. F. Microbial dysbiosis-induced obesity: role of gut microbiota in homoeostasis of energy metabolism. Br. J. Nutr. 123, 1127–1137 (2020).

47. Kavanagh, P. et al. Tentative identification of the phase I and II metabolites of two synthetic cathinones, MDPHP and α-PBP, in human urine. Drug Test. Anal. 12, 1442–1451 (2020).

48. Wang, J.-H. et al. Clinical evidence of the link between gut microbiome and myalgic encephalomyelitis/chronic fatigue syndrome: a retrospective review. Eur. J. Med. Res. 2G, 148 (2024).

49. Lenoir, M. et al. Butyrate mediates anti-inflammatory effects of Faecalibacterium prausnitzii in intestinal epithelial cells through Dact3. Gut Microbes (2020).

50. Sokol, H. et al. Faecalibacterium prausnitzii is an anti-inflammatory commensal bacterium identified by gut microbiota analysis of Crohn disease patients. Proc. Natl. Acad. Sci. 105, 16731–16736 (2008).

51. Quévrain, E., et al. Identification of an anti-inflammatory protein from Faecalibacterium prausnitzii, a commensal bacterium deficient in Crohn’s disease. Gut 65, 415–425 (2016).

52. Miquel, S. et al. Identification of Metabolic Signatures Linked to Anti-Inflammatory Effects of Faecalibacterium prausnitzii. mBio 6, 10.1128/mbio.00300-15 (2015).

53. Vital, M., Howe, A. C. C, Tiedje, J. M. Revealing the Bacterial Butyrate Synthesis Pathways by Analyzing (Meta)genomic Data. mBio 5, e00889–14 (2021).

54. Recharla, N., Geesala, R. C, Shi, X.-Z. Gut Microbial Metabolite Butyrate and Its Therapeutic Role in Inflammatory Bowel Disease: A Literature Review. Nutrients 15, 2275 (2023).

55. Monteiro, C. R. A. V., et al. In Vitro Antimicrobial Activity and Probiotic Potential of Bifidobacterium and Lactobacillus against Species of Clostridium. Nutrients 11, 448 (2019).

56. Zhao, M., Li, G. C, Deng, Y. Engineering Escherichia coli for Glutarate Production as the C5 Platform Backbone. Appl. Environ. Microbiol. 84, e00814–18 (2018).

57. Nguyen-Lefebvre, A. T., Selzner, N., Wrana, J. L. C, Bhat, M. The hippo pathway: A master regulator of liver metabolism, regeneration, and disease. FASEB J. 35, e21570 (2021).

58. Khan, M. A., Gupta, A., Sastry, J. L. N. C, Ahmad, S. Hepatoprotective potential of kumaryasava and its concentrate against CCl4-induced hepatic toxicity in Wistar rats. J. Pharm. Bioallied Sci. 7, 297–299 (2015).

59. Kim, C.-S. Roles of Diet-Associated Gut Microbial Metabolites on Brain Health: Cell-to-Cell Interactions between Gut Bacteria and the Central Nervous System. Adv. Nutr. 15, 100136 (2024).

60. Rebeaud, J., Peter, B. C, Pot, C. How Microbiota-Derived Metabolites Link the Gut to the Brain during Neuroinflammation. Int. J. Mol. Sci. 23, 10128 (2022).

61. Ahmad, S. et al. Gut microbiome-related metabolites in plasma are associated with general cognition. Alzheimers Dement. 17, e056142 (2021).

62. Ahmed, Z., Zeeshan, S., Xiong, R. C, Liang, B. T. Debutant iOS app and gene-disease complexities in clinical genomics and precision medicine. Clin. Transl. Med. 8, e26 (2019).

63. Ahmed, Z., Zeeshan, S., Xiong, R. C, Liang, B. T. PAS-Gen: Guide to iOS app with gene-disease.

64. Ahmed, Z., Wan, S., Zhang, F. C, Zhong, W. Artificial intelligence for omics data analysis. BMC Methods 1, 4 (2024).

65. Ahmed, Z., Mohamed, K., Zeeshan, S. C, Dong, X. Artificial intelligence with multi-functional machine learning platform development for better healthcare and precision medicine. Database J. Biol. Databases Curation 2020, baaa010 (2020).

66. Xiong, R. et al. Multi-‘omics of gut microbiome-host interactions in short- and long-term myalgic encephalomyelitis/chronic fatigue syndrome patients. Cell Host Microbe 31, 273–287.e5 (2023).

67. Germain, A., Ruppert, D., Levine, S. M. C, Hanson, M. R. Prospective Biomarkers from Plasma Metabolomics of Myalgic Encephalomyelitis/Chronic Fatigue Syndrome Implicate Redox Imbalance in Disease Symptomatology. Metabolites 8, 90 (2018).

68. Lim, E.-J. C, Son, C.-G. Review of case definitions for myalgic encephalomyelitis/chronic fatigue syndrome (ME/CFS). J. Transl. Med. 18, 289 (2020).

69. Germain, A., Barupal, D. K., Levine, S. M. C, Hanson, M. R. Comprehensive Circulatory Metabolomics in ME/CFS Reveals Disrupted Metabolism of Acyl Lipids and Steroids. Metabolites 10, 34 (2020).

70. Jason, L. A., Yoo, S. C, Bhatia, S. Patient perceptions of infectious illnesses preceding Myalgic Encephalomyelitis/Chronic Fatigue Syndrome. Chronic Illn. 18, 901–910 (2022).

71. Hanson, M. R. The viral origin of myalgic encephalomyelitis/chronic fatigue syndrome. PLOS Pathog. 19, e1011523 (2023).

72. Hamine, S., Gerth-Guyette, E., Faulx, D., Green, B. B. C, Ginsburg, A. S. Impact of mHealth Chronic Disease Management on Treatment Adherence and Patient Outcomes: A Systematic Review. J. Med. Internet Res. 17, e3951 (2015).

73. Clark, N. M. Management of Chronic Disease by Patients. Annu. Rev. Public Health 24, 289–313 (2003).

74. Derman, I. D. et al. High-throughput bioprinting of the nasal epithelium using patient-derived nasal epithelial cells. Biofabrication 15, 044103 (2023).

75. Fleming, E. et al. Cultivation of common bacterial species and strains from human skin, oral, and gut microbiota. BMC Microbiol. 21, 278 (2021).

76. Vyas, J., Muirhead, N., Singh, R., Ephgrave, R. C, Finlay, A. Y. Impact of myalgic encephalomyelitis/chronic fatigue syndrome (ME/CFS) on the quality of life of people with ME/CFS and their partners and family members: an online cross-sectional survey. BMJ Open 12, e058128 (2022).

77. Martinez, A., Okoh, A., Ko, Y.-A. C, Wells, B. Racial Differences in FMD. 2023.02.10.23285630 Preprint at 10.1101/2023.02.10.23285630 (2023).

78. Trivedi, M. S. et al. Identification of Myalgic Encephalomyelitis/Chronic Fatigue Syndrome-associated DNA methylation patterns. PLOS ONE 13, e0201066 (2018).

79. Bouquet, J. et al. Whole blood human transcriptome and virome analysis of ME/CFS patients experiencing post-exertional malaise following cardiopulmonary exercise testing. PLOS ONE 14, e0212193 (2019).

80. Lande, A. et al. Human Leukocyte Antigen alleles associated with Myalgic Encephalomyelitis/Chronic Fatigue Syndrome (ME/CFS). Sci. Rep. 10, 5267 (2020).

81. Almenar-Pérez, E. et al. Epigenetic Components of Myalgic Encephalomyelitis/Chronic Fatigue Syndrome Uncover Potential Transposable Element Activation. Clin. Ther. 41, 675–698 (2019).

82. Das, S., Taylor, K., Kozubek, J., Sardell, J. C, Gardner, S. Genetic risk factors for ME/CFS identified using combinatorial analysis. J. Transl. Med. 20, 598 (2022).

83. Caruana, E. J., Roman, M., Hernández-Sánchez, J. C, Solli, P. Longitudinal studies. J. Thorac. Dis. 7, E537–E540 (2015).

84. White, R. T. C, Arzi, H. J. Longitudinal Studies: Designs, Validity, Practicality, and Value. Res. Sci. Educ. 35, 137–149 (2005).

85. Aurora, C., Cecilia, A. C, Adina, H. The Role of Diet in the Treatment of Chronic Diseases Case Study. ARS Medica Tomitana 27, 153–156 (2021).

86. Therrien, R. C, Doyle, S. Role of training data variability on classifier performance and generalizability. in Medical Imaging 2018: Digital Pathology vol. 10581 58–70 (SPIE, 2018).

87. Zhang, B., Qin, A. K., Pan, H. C, Sellis, T. A Novel DNN Training Framework via Data Sampling and Multi-Task Optimization. in 2020 International Joint Conference on Neural Networks (IJCNN) 1–8 (2020). doi:10.1109/IJCNN48605.2020.9207329.

88. Krumina, A. et al. Clinical Profile and Aspects of Differential Diagnosis in Patients with ME/CFS from Latvia. Medicina (Mex.) 57, 958 (2021).

89. Zubcevik, N. et al. Symptom Clusters and Functional Impairment in Individuals Treated for Lyme Borreliosis. Front. Med. 7, (2020).

90. Costa, G. G., Pereira, A. R. C, Carvalho, A. S. Pericardite lúpica: dor torácica e febre em tempos de COVID-19. Rev. Port. Med. Geral E Fam. 38, 300–4 (2022).

91. Lathan, C., Spira, J. L., Bleiberg, J., Vice, J. C, Tsao, J. W. Defense Automated Neurobehavioral Assessment (DANA)-psychometric properties of a new field-deployable neurocognitive assessment tool. Mil. Med. 178, 365–371 (2013).

92. Resnick, H. E. C, Lathan, C. E. From battlefield to home: a mobile platform for assessing brain health. mHealth 2, 30 (2016).

93. Lee, J. et al. Hemodynamics during the 10-minute NASA Lean Test: evidence of circulatory decompensation in a subset of ME/CFS patients. J. Transl. Med. 18, 314 (2020).

94. Committee on the Diagnostic Criteria for Myalgic Encephalomyelitis/ChronicFatigue Syndrome, Board on the Health of Select Populations, & Institute of Medicine. Beyond Myalgic Encephalomyelitis/Chronic Fatigue Syndrome: Redefining an Illness. (National Academies Press (US), Washington (DC), 2015).

95. Monica, 1776 Main Street Santa & California 90401-3208. 36-Item Short Form Survey (SF-36) Scoring Instructions. https://www.rand.org/health-care/surveys_tools/mos/36-item-short-form/scoring.html.

96. Shen, W., Le, S., Li, Y. C, Hu, F. SeqKit: A Cross-Platform and Ultrafast Toolkit for FASTA/Q File Manipulation. PLOS ONE 11, e0163962 (2016).

97. Blanco-Míguez, A. et al. Extending and improving metagenomic taxonomic profiling with uncharacterized species using MetaPhlAn 4. Nat. Biotechnol. 41, 1633–1644 (2023).

98. Edgar, R. C. Search and clustering orders of magnitude faster than BLAST. Bioinformatics 26, 2460–2461 (2010).

99. Love, M. I., Huber, W. C, Anders, S. Moderated estimation of fold change and dispersion for RNA-seq data with DESeq2. Genome Biol. 15, 550 (2014).

100. Mallick, H. et al. Multivariable association discovery in population-scale meta-omics studies. PLoS Comput. Biol. 17, e1009442 (2021).

101. Chawla, N. V., Bowyer, K. W., Hall, L. O. C, Kegelmeyer, W. P. SMOTE: Synthetic Minority Over-sampling Technique. J. Artif. Intell. Res. 16, 321–357 (2002).

102. Haibo He, Yang Bai, Garcia, E. A., & Shutao Li. ADASYN: Adaptive synthetic sampling approach for imbalanced learning. in 2008 IEEE International Joint Conference on Neural Networks (IEEE World Congress on Computational Intelligence) 1322–1328 (IEEE, Hong Kong, China, 2008). doi:10.1109/IJCNN.2008.4633969.

103. Saripuddin, M., Suliman, A., Syarmila Sameon, S. & Jorgensen, B. N. Random Undersampling on Imbalance Time Series Data for Anomaly Detection. in *Proceedings of the 2021 4th International Conference on Machine Learning and Machine Intelligence* 151– 156 (Association for Computing Machinery, New York, NY, USA, 2022). doi:10.1145/3490725.3490748.

104. Lundberg, S. C, Lee, S.-I. A Unified Approach to Interpreting Model Predictions. Preprint at 10.48550/arXiv.1705.07874 (2017).

105. Abadi, M., et al. TensorFlow: A system for large-scale machine learning. Preprint at 10.48550/arXiv.1605.08695 (2016).

106. Pedregosa, F., et al. Scikit-learn: Machine Learning in Python. Preprint at 10.48550/arXiv.1201.0490 (2018).

107. Friedman, J. H., Hastie, T. C, Tibshirani, R. Regularization Paths for Generalized Linear Models via Coordinate Descent. J. Stat. Softw. 33, 1–22 (2010).

108. Kuhn, M. Building Predictive Models in R Using the caret Package. J. Stat. Softw. 28, 1–26 (2008).

109. Langfelder, P. C, Horvath, S. WGCNA: an R package for weighted correlation network analysis. BMC Bioinformatics 9, 559 (2008).

110. Antonov, M., et al. igraph enables fast and robust network analysis across programming languages. Preprint at 10.48550/arXiv.2311.10260 (2023).

111. Koo, T. K. C, Li, M. Y. A Guideline of Selecting and Reporting Intraclass Correlation Coefficients for Reliability Research. J. Chiropr. Med. 15, 155–163 (2016).

112. Nelder, J. A. C, Wedderburn, R. W. M. Generalized Linear Models. J. R. Stat. Soc. Ser. Gen. 135, 370–384 (1972).

113. Bakdash, J. Z. C, Marusich, L. R. Repeated Measures Correlation. Front. Psychol. 8, (2017).

114. Gu, C. Smoothing Spline ANOVA Models: R Package gss. Smoothing Spline ANOVA Models.

